# Ion Channel Nano-Diagnostics for ER+ Breast Cancer

**DOI:** 10.64898/2026.03.09.710404

**Authors:** Manos Gkikas, Evangelos Dadiotis, Mehreen Zaka, Nihal Aly, Kim Chan, Diomedes. E. Logothetis

**Affiliations:** Northeastern University, Department of Pharmaceutical Sciences, Boston, Massachusetts 02115, U.S.A

## Abstract

Ion channels are pore-forming transmembrane proteins that allow ions to move down an electrochemical gradient and across the channel pore and regulate many cell functions. Among them, are the G-protein–gated inwardly-rectifying K+ channels 1 (GIRK1) that are ubiquitously expressed with major functions in the brain and heart. Interestingly, significantly higher GIRK1 expression has been found in estrogen receptor positive (ER+) breast cancer patients compared to patients with HER2+ tumors or normal patients, and that was statistically correlated with shorter survival times and metastatic potential. Herein, we report the preparation of ∼4 nm *GAT1508*-coated poly(ethylene glycol) gold nanoparticle (PEGylated AuNP) biomarker for ER+ breast cancer cell screening through an optical microscope. A urea-based small molecule, GAT1508, with an N-methylpyrazole benzyl group on one side and a bromo-thiophene tail on the other side, has been shown to predominantly bind GIRK1 subunits and specifically activate GIRK1/2 channels. Two derivatives of GAT1508were synthesized and characterized: an ethylamine derivative (GAT1508-EA) with a chain extension from the benzyl ring, and a propylamine derivative (GAT1508-PA) with a chain extension from the pyrazole ring. Electrophysiology (TEVC and whole-cell patch-clump) experiments as well as fluorescence studies (Thallium assay) showed that only GAT1508-PA inhibited GIRK1/2-mediated K+ currents in transfected HEK293^GIRK1^ cells. Docking studies showed strong binding for the propylamine GAT1508 derivative, both in the amine form (GAT1508-PA) as well as in the amide form (GAT1508-PA-EG2; coupled with PEG as in the AuNPs). *GAT1508*-PEG-AuNPs (*GAT1508*-NPs) were synthesized subsequently with ∼65 wt% metal loading. UV-Vis studies revealed the presence of the conjugated ligand at 260 nm. Flow cytometry studies showed binding of Alexa 594-labeled *GAT1508*-NPs in ER+ MCF-7 breast cancer cells with a strong interaction, while incubation of fixed MCF-7 cells with a *GAT1508*-NP solution led to optical detection of ER+ breast cancer cells, without the need of fluorescent dyes and additional amplification steps. Detection was not feasible in MDA-MB-231 cells, a triple (-) breast cell line that does not express GIRK1. This is the first study, to our knowledge, that couples nanotechnology with small molecule drug design and electrophysiology to develop ion channel-tracing molecular probes for the detection/screening of ER+ breast cancer.

## Introduction

Breast cancer is the most common cancer in women in the United States,^1–5^ accounting for ∼30% of all new female cancers, according to the American Cancer Society. Each year, around half a million people are diagnosed with breast cancer and approximately 1/5 of women decease,^5^ while the number is expected to increase by 39% in 2040. Breast cancer screening is known to reduce the mortality rates by 20%.^6^ Medical imaging, hematoxylin and eosin (H&E) staining, immunohistochemistry, mRNA quantification, and RNA in situ hybridization (ISH)^7^ are used for tissue diagnostics (histology), apart from tumor cell smears (cytology), which are rarely used for breast cancer diagnosis unlike other cancers. Even though all these methods work effectively, they require significant time to report results (∼7-10 days for immunohistochemistry), or cost, demanding a high number of reagents to develop (e.g. different tissue sections must be pretreated for RNA-ISH, then stained with 3 different probes, followed by x6 amplification steps, and an extra reaction with DAB: 3,3′-diaminobenzidine for signal detection). For a hospital that receives a high number of breast cancer samples per day this is challenging and can become even more difficult in cases of short-staffed personnel. The time required for patients to receive biopsy/ tissue diagnosis, as well as the cost (both immunohistochemistry and RNA-ISH), motivates the development of a rapid and economical diagnostic method for early breast cancer detection.

Potassium ion channels are dysregulated in breast cancer due to a number of factors that collectively enable cancer cells to control their membrane potential and signaling to favor proliferation, migration, and resistance to cell death.^8^ G-protein–gated inwardly-rectifying K+ (GIRK) channels have recently received increased attention in breast (∼40% of breast cancer tissues shave shown high mRNA expression^9^) and metastatic cancer due to their high expression in those cells.^7,9–11^ GIRK channels can be found in the cell membrane as functional homotetramers (GIRK2 and GIRK4) or heterotetramers (e.g., GIRK1/2, GIRK1/4, GIRK2/3).^12^ Stringer et al. found increased levels of KCNJ3 (the gene that encodes GIRK1) mRNA in primary breast carcinomas compared to normal breast tissue, and noticed a positive correlation between KCNJ3 levels in the tumor and the number of metastatic lymph nodes. ^10^ By examining a high number of clinical samples, the groups of Schreibmayer and Bauernhofer found significantly higher GIRK1 expression levels in patients with ER+ breast cancer (both luminal A and B subtypes) compared to patients with HER2+ tumors or normal patients.^7^ Interestingly, comparison of >1000 breast cancer patients grouped solely based on ER status revealed substantially higher KCNJ3 levels in ER+ vs. ER- patients (p<0.001).^7^ Statistical analysis showed correlation of KCNJ3 expression with ERα (p<0.001; found mainly in breast and uterus) but not with ERβ (mainly found in the ovaries and brain) or the G protein-coupled ER, while survival analysis of ER+ patients indicated that *patients with high GIRK1 levels in the tumor had also shorter overall survival times* than patients with low KCNJ3 levels (n = 647 patients; p<0.05).^7^

Since ER+ breast cancer patients can be classified into *low* and *high risk* groups based on the GIRK1 levels in the tumor (independent prognostic marker),^7^ developing a rapid method to detect and amplify the presence of overexpressed GIRK1 channels in cells/tissues could be of paramount importance for clinical evaluation of ER+ breast cancer. This could fill the existing gap of absence of anti-GIRK1 antibodies for immunohistochemistry since several of those tested did not meet the required high quality sensitivity and specificity standards.^7^ Nanotechnology could play a crucial role towards that direction.^13–23^ GAT1508 was recently introduced as a selective ligand (activator) for GIRK1/2 channels in the brain, without affecting GIRK1/4 channels in the heart, unlike the commercially available ligand ML297.^12^ It was envisioned that a potential modification of GAT1508 could allow its conjugation to NPs, and combine the enhanced light-absorbance of small-size NPs with the strong binding of the ligand to the overexpressed GIRK1 channels in breast cancer cells, enabling anchoring of NPs to cancerous cells/tissues and tracing of ER+ breast cancer through the upregulated transmembrane GIRK1 channels.

Herein, we report a novel screening method for ER+ breast cancer through the detection of the overexpressed GIRK1 channels in ER+ breast cancer cells via ligand-bound NPs. Two GAT1508 alkylamine derivatives were first synthesized, and their binding to GIRK1/2 channels was examined using three different methods (TEVC: two-electrode voltage clamp, MPC: manual whole-cell patch clamp, and TFA: Thallium fluorescence assay). Docking studies and Molecular Dynamic (MD) simulations revealed strong affinity interactions between the ligand and the receptor. The most promising ligand was then conjugated to PEGylated AuNPs yielding ∼4 nm *GAT1508*-PEG-AuNPs. Flow cytometry revealed binding of the NPs to ER+ MCF-7 cells. Incubation of fixed MCF-7 cells using a *GAT1508*-NP solution enabled optical microscope visualization of ER+ breast cancer cells through the bound NPs to the overexpressed GIRK1 channels, simplifying the detection method for broad hospital screening applications. On the contrary, no detection was found when a triple (-) human breast cell line was used that did not express GIRK1. Having shown that it works in cells, we envision testing our technology with cancerous tissues. If successful, our technology could reduce by 1/20 the time required for clinical ER+ tumor diagnosis in histology slices, minimizing the waiting time for patients, lowering the diagnostic test cost, and giving accessibility to low income groups, female veterans and on duty personnel, enabling rapid and low-cost breast cancer diagnostics that can cover rural areas in the US and Europe as well as parts in Africa where only breast palpation is currently used for diagnosis.

## Results and Discussion

### Synthesis of GAT1508 Alkylamine Derivatives (GAT1508-EA and GAT1508-PA)

GAT1508 selectively activates GIRK1/2 channels and allows K^+^ ion permeation.^12^ However, this molecule cannot be used in its current form for incorporation into nanomaterials, as it lacks a functional group that can serve as an anchoring site for surface attachment. Alkylamine derivatives of GAT1508 extending either from the benzyl ring (GAT1508 ethylamine: GAT1508-EA) or the pyrazole ring (GAT1508 propylamine: GAT1508-PA) were thus synthesized (chemical structures in **Figure 1a**) following a 5 min microwave-catalyzed reaction at 120 °C between 5-bromothiophene-2-carboxylic acid and benzyl pyrazole alkylated amines (**Scheme S1**). Higher yield was achieved when the reactant amines were in the salt form vs. free amine. The purity of the synthesized products was verified by ^1^H-NMR and MS spectroscopies (**Figures S1**-**S4**). GAT1508-EA (extending from the benzyl ring) showed no interaction with GIRK1/2 channels overexpressed in HEK293 cells (**Figure S5**). However, GAT1508-PA (extending from the pyrazole ring) showed partial inhibition of GIRK1/2-mediated K+ currents in transfected oocytes and HEK293 cells and strong binding with the GIRK1 channels, as shown by three methods: TEVC, MPC, and the TFA. TEVC showed partial inhibition of the high K^+^ (HK) solution using 10 μM of GAT1508-PA (**Figure 1b**), while MPC showed a similar effect with 30 μM of the same ligand (**Figure 1c**), with statistical significance from the high K^+^ (HK) solution (**Figure 1d**). The effect was evident even in the coexistence of 5 μM of GAT1508 (channel activator) with 30 μM of GAT1508-PA (**Figure 1e**), showing ∼25% inhibition of the activated channel. TFA, which exploits the permeation of thallium (Tl) ions via ligand-gated K^+^ channels (GIRK1/2 in our case) through binding to a highly-sensitive fluorescent dye inside the cell yielding strong fluorescence in the case of GAT1508, led to a dose-dependent reduction in the Tl^+^/dye signal by GAT1508-PA (**Figure 1f**). The inhibitory effect of the synthesized GAT1508-PA derivative denotes binding to GIRK1 channels. This is also confirmed by computational docking studies (see below). For an *ex vivo* application, such as the breast cancer detection in tissues, the ligand that binds to the GIRK1 channels can be either an activator or an inhibitor, providing anchoring of the contrasting molecule to the overexpressed ion channels of breast cancer cells/tumors.

**Figure 1.**
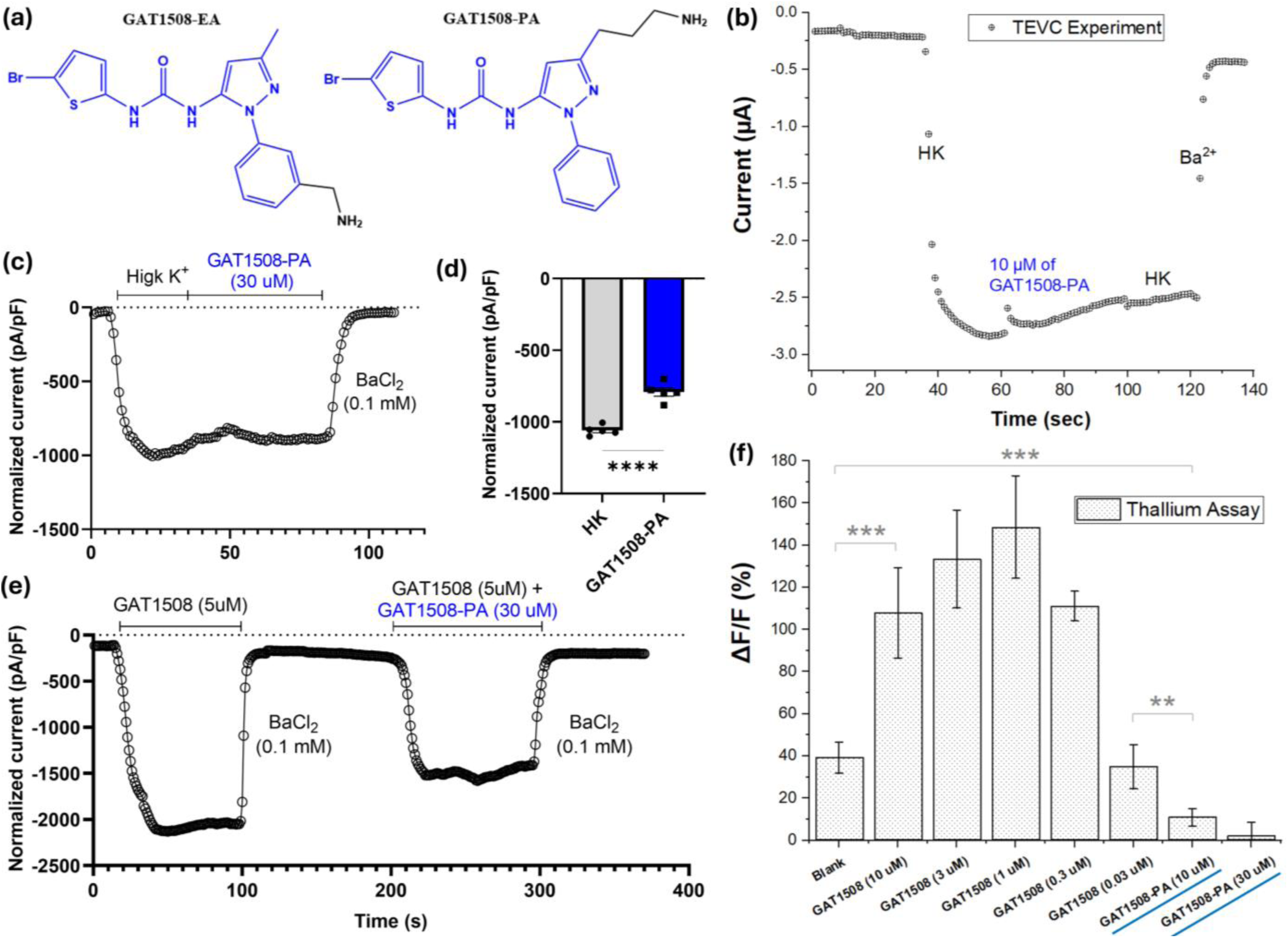
(a) Chemical structures of the alkylamine derivatives GAT1508-EA and GAT1508-PA; (b) TEVC of transfected HEK293T^GIRK1/2^ cells in the presence of high K^+^ solution (natural channel activator), 10 μM GAT1508-PA, and Ba^2+^ (ion channel blocker); (c) MPC (whole-cell) recordings of HEK293^GIRK1/2^ cells with high K^+^ solution and 30 μM of GAT1508-PA, resulting in (d) statistically significant inhibitory effect on GIRK1/2-mediated currents compared to high K^+^ solution (data are presented as mean ± SEM, p < 0.0001, unpaired t-test); (e) current response of HEK293^GIRK1/2^ cells using 5 μM GAT1508 alone and in the presence of 30 μM of GAT1508-PA; (f) TFA showing increase in fluorescence (Tl^+^/dye complex) for GAT1508 by itself (10 - 0.3 μΜ) and signal reduction dose-dependently on GAT1508-PA (10 μΜ and 30 μM).

### Docking of GAT1508-PA Derivatives to GIRK1/2 Channels

GIRK channels have been broadly studied by the Logothetis lab^12,23–33^ and are regulated by phosphatidylinositol 4,5-bisphosphate (PIP2). PIP₂ plays a pivotal role in ion channel gating and ligand modulation, and its presence is essential for channel activation or stabilization of specific conformational states. Incorporating PIP₂ into our computational models, particularly in gating molecular dynamics (GMD) allows for more physiologically relevant predictions of ligand binding. Experimental and model predictions highlight the importance of GIRK1 in its ability to form polar interactions with the PIP2 phosphates to complement corresponding interactions with the GIRK2 or GIRK4 subunits.^34,35^

The binding affinity of the GAT1508-PA was explored by docking studies. Previous studies have shown that GAT1508 binds mainly to the GIRK1 chain through interactions (hydrogen bonding) of the *urea group* with amino acids Asp-173 and Trp-95.^12^ Cryo-EM structure of GIRK1/2 channels with GAT1508 revealed that its *bromothiophene* site is engaged in a π-network (T-stacking) with Met-98.^36^ GAT1508-PA was found to bind GIRK1/2 channels with high affinity, defined by docking score and MMGBSA dG_bind_ score (Molecular Mechanics/Generalized Born Surface Area; Table in **Figure 2a**). GMD also confirmed that GAT1508-PA has low K^+^ ion permeation depicting the inhibitory effect of the compound. **Figure 2b** shows the 2D interaction diagram of GAT1508-PA highlighting the key residues Asp-173 and Trp-95 making hydrogen bond interactions with the urea group of the compound, and keeping the ligand stably in the binding pocket. This binding pose was stable during the 500 ns of molecular dynamics simulations. GMD was performed in 3 replicas to confirm the overall binding affinity and inhibitory nature of the compound. Docking and MMGBSA studies were also performed for the GAT1508-PA derivative with oligo(ethylene glycol) (GAT1508-PA-EG2) to mimic the actual structure of the ligand conjugated to the NP via amide bond through a PEG linker. Overall, both GAT1508-PA and GAT1508-PA-EG2 are stable in the binding pocket of GIRK1/2 and do not activate the channel in the time frame of the simulations, in contrast with GAT1508. The 2D interaction diagram of GAT1508 and GAT1508-EG2 shows that Asp-173 and Trp-95 are the crucial residue interactions for keeping the ligands stable in the binding pocket (**Figure S6**).

**Figure 2.**
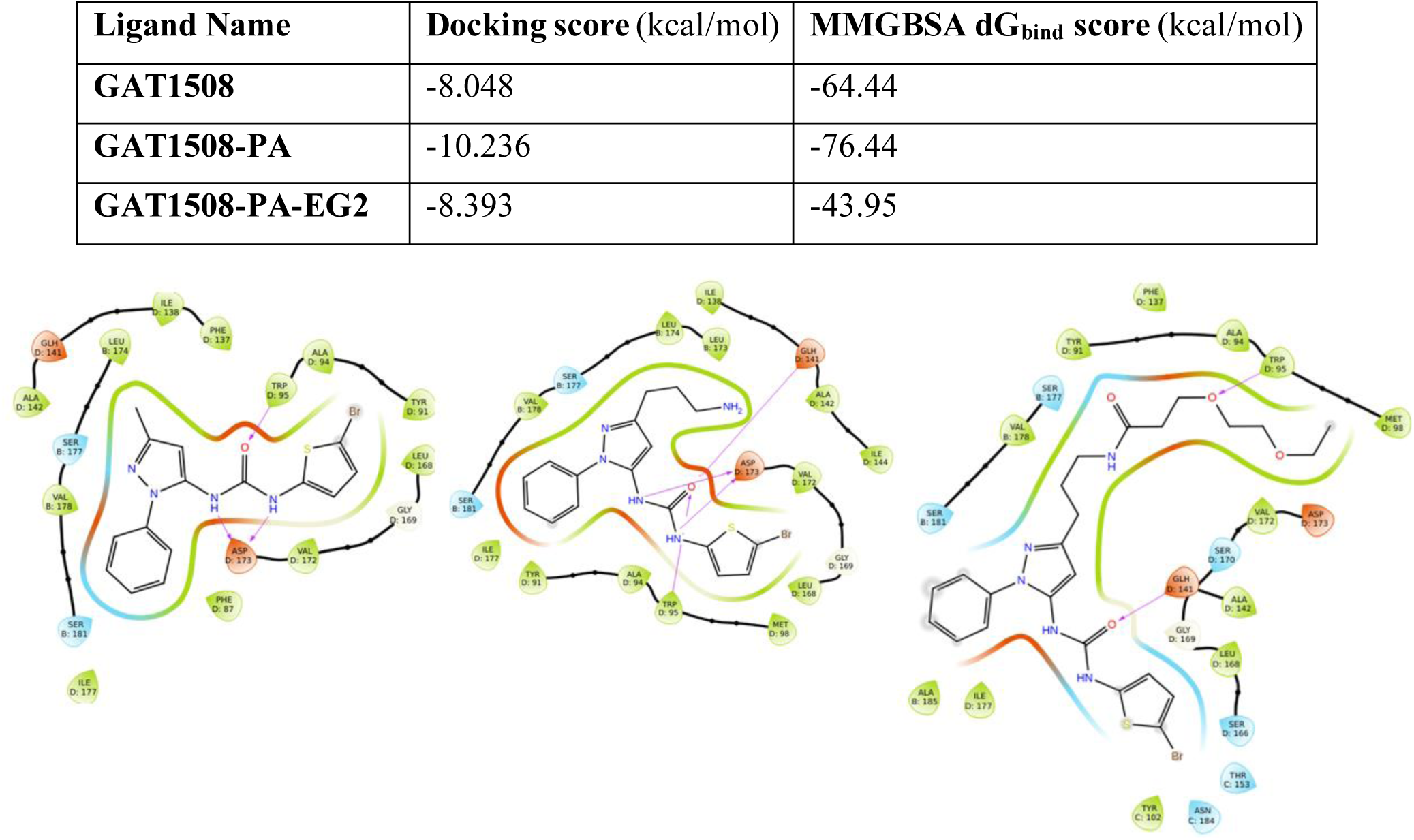
(a) Table containing docking and MMGBSA dG_bind_ scores of GAT1508, GAT1508-PA, and GAT1508-PA-EG2; (b) 2D interactions of the ligands above in the binding pocket of GIRK1/2 channel.

### Synthesis of HOOC-PEG-Coated Gold Nanoparticles (HOOC-PEG-AuNPs)

A matrix-free^15–17,19^ hydrophilic contrast probe precursor, HOOC-PEG-AuNPs, was synthesized by ligand exchange of 3.7 ± 0.6 nm (**Figure S7**) dodecanethiol-coated AuNPs (∼**11.7** ± 0.3 **wt%** dodecanethiol incorporation by TGA: thermogravimetric analysis; decomposition temperature: *T_d_ = 301 ± 5 °C*; **Figure S8**) with excess HOOC-PEG_5K_-SH (PEG had a **96.6** ± 0.2 **wt%** loss with *T_d_*= *416 ± 1 ℃*; TGA in **Figure S9**) followed by precipitation in hexane to remove liberated dodecanethiol. PEG improves the NP solubility/dispersibility due to hydration of the NP core, reduces non-specific protein adsorption, provides stealth properties in the bloodstream due to its dysopsonic properties, and prolongs the half-life of PEG-conjugated materials.^37–39^ High speed centrifugation (fractionation) was then used to remove unbound HOOC-PEG-SH, since free polymeric chains have been found to co-exist with polymer-coated NP in previous studies even after dialysis.^15–17,19^ Post high-speed fractionation, the purified HOOC-PEG-AuNPs showed two weight losses; one with **9.1** ± 0.2 **wt%** loss (*T_d_* = *278 ± 3 ℃*) due to the decomposition of remaining dodecathiol, and another one between 270-520℃ with **34.5** ± 0.8 **wt%** loss (*T_d_* = *401 ± 3 ℃*) due to the decomposition of NP-bound PEG-SH (**Figure 3a**; all TGAs are shown analytically in **Figure S10**). Unlike the hydrophobic C12-S-AuNPs which are dispersible in toluene, HOOC-PEG-AuNPs were highly dispersible in water. The average size of uncoated PEGylated AuNPs was **3.2** ± 0.4 **nm** (**Figure S11**).

**Figure 3.**
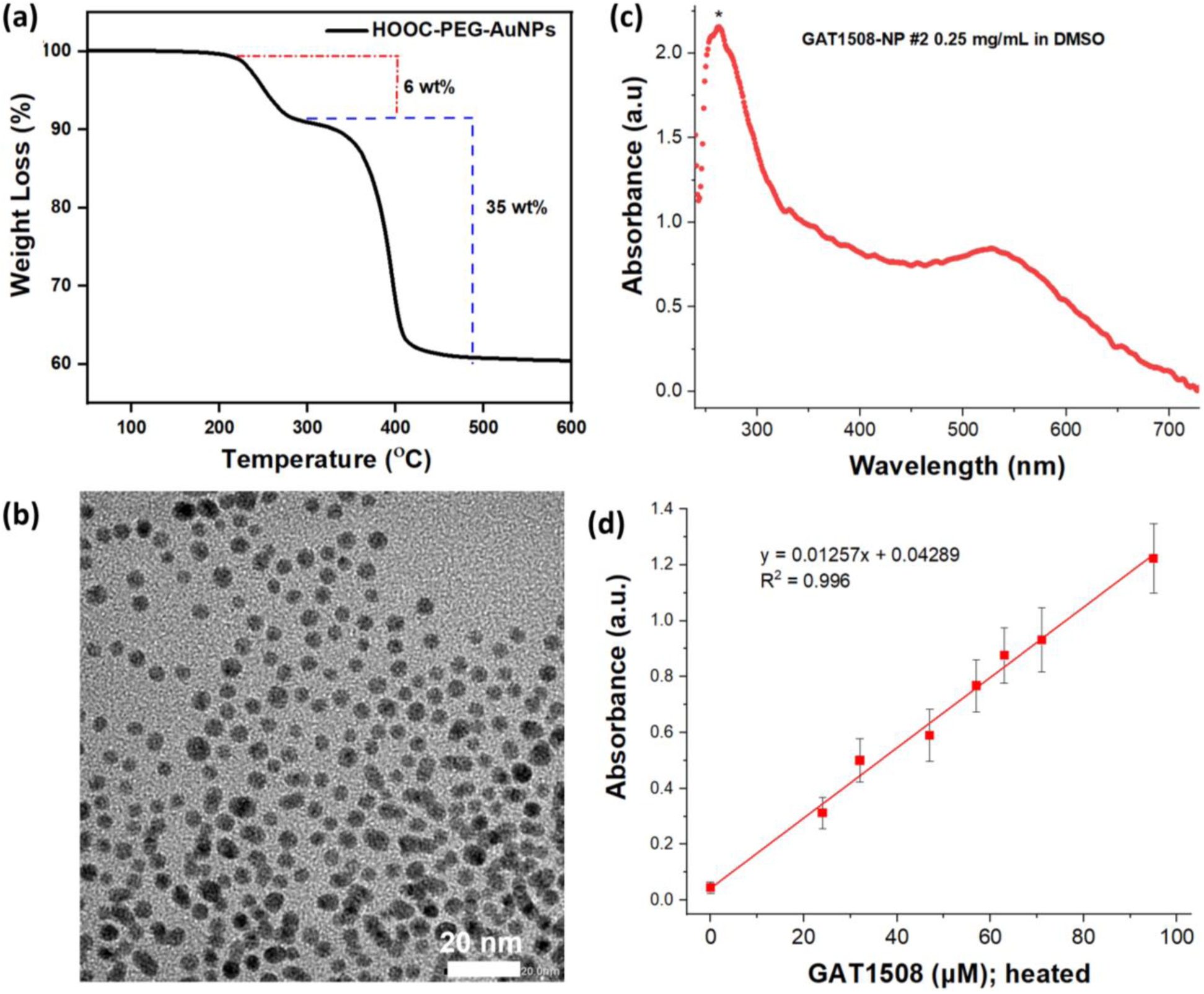
(a) TGA curve of HOOC-PEG-AuNPs; (b) TEM image of *GAT1508*-PEG-AuNPs; (c) UV-Vis spectrum of *GAT1508*-PEG-AuNPs at 0.25 mg/mL in DMSO. The conjugated ligand peak at 280 nm is shown with an asterisk; (d) Linear fitting of the UV-Vis peak of *GAT1508* at 261 nm between 0-95 μM in DMSO. The equation was used to quantify the conjugated *GAT1508* after hydrolysis of the amide bond in NP-ligand. The absorbance of ligand-free NP was subtracted.

### Synthesis of *GAT1508*-Coated PEG-Gold Nanoparticles (*GAT1508*-PEG-AuNPs)

Matrix-free HOOC-PEG-AuNPs were subsequently conjugated with GAT1508-PA via EDC/NHS-catalyzed amide coupling (overnight reaction) followed by extensive dialysis (3-4 days) over DI water to remove unbound drug, and lyophilized. Purified *GAT1508*-PEG-AuNPs had an average particle size of **4.2** ± 0.6 **nm** (**Figure 3b**; analytically in **Figure S12**). The NP size in solution was examined by UV-Vis spectroscopy. HOOC-PEG-AuNPs showed a surface plasmon resonance (SPR) band at 520 nm, whereas *GAT1508*-PEG-AuNPs had a SPR band at 526 nm (**Figure 3c**). The latter, in addition, had an absorption peak at 260 nm, which is indicative of GAT1508 (see calibration curve in **Figure S13**). To quantify the NP-bound ligand, *GAT1508*-PEG-AuNPs were hydrolyzed in 6 N HCl at 105 °C for 24 h, the solvent was evaporated, and the sample was dissolved in 1 mL DMSO and quantified by UV-Vis after 5 min centrifugation (at 16,000 rpm) to isolate the supernatant which contained the DMSO-soluble GAT1508 from the solid NPs which remained in the pellet. A calibration curve of the ligand, processed at the same conditions, was used as a reference (**Figure 3d**). It was found that 2 mg of GAT1508-NPs/mL contained 79 μM of *GAT1508* after absorbance correction over ligand-free NP, or 39.5 μM *GAT1508*/mg NP.

### Optical Microscope Detection of ER+ MCF-7 Cells Using *GAT1508*-PEG-AuNPs

Histology samples of ER+ breast cancer patients have shown high levels of GIRK1 levels based on mRNA quantification.^7^ Choosing a representative ER+/GIRK1+ cell line is thus highly important for cell studies. According to the *Human Protein Atlas*, human ER+ MCF-7 breast cancer cells have the highest levels of GIRK1, as well as a small number of GIRK2 channels, while GIRK4 channels are absent. GIRK1 alone does not form functional homotetramers and instead remains as a cytosolic (endoplasmic reticulum: ER-trapped) protein.^40,41^ However, the presence of a small amount of GIRK2, even at RNA ratios as high as 20:1 GIRK1/GIRK2 (4 + 0.2), results in robust channel activity as shown in the *Xenopus* oocyte heterologous expression system (**Figure S14**). This indicates that even a relatively small presence of GIRK2 is sufficient to promote the assembly of GIRK1 into transmembrane tetrameric units, enabling the formation of functional channels.

*GAT1508*-NPs (with GAT1508-PA ligand incorporated; **Figure 4a**) showed strong interaction with fixed, ER+ MCF-7 breast cancer cells (**Figure 4b,c**) post incubation into a NP dispersion, evident by enhanced Vis contrast, most probably due to interaction with the overexpressed GIRK1/2 channels. The effect was detected after x2 washings with PBS to remove unbound NPs, and was evident throughout the sample (highlighted in green circles in **Figure 4b**) to denote the statistics. Small-size NPs (∼4 nm in our case) are known to absorb light (peak max at 526 nm) stronger than scattering light, which occurs with larger size NPs.^42^ It is important to note that our nanotechnology worked with ER+ breast cancer cells adherent both onto cell culture flasks as well as onto cover slips. In addition, the *GAT1508*-NP solution managed to give reproducible results even after 2 weeks of storage (not examined further), and allowed for the development of multiple ER+ breast cancer cell samples grown and fixed in cover slips using the same *GAT1508*-NP solution. Images on a different (basic) optical microscope were also obtained using a cell phone camera (**Figure S15**) to show the applicability of our method. A high-resolution cryo-EM structure of GIRK1/2 binding to GAT1508 has been obtained.^36^ On the contrary, absence of binding was achieved when *GAT1508-*NPs were incubated with triple (-) MDA-MB-231 cells (**Figure S16**), a control cell line that does not express ER and GIRK1.

**Figure 4.**
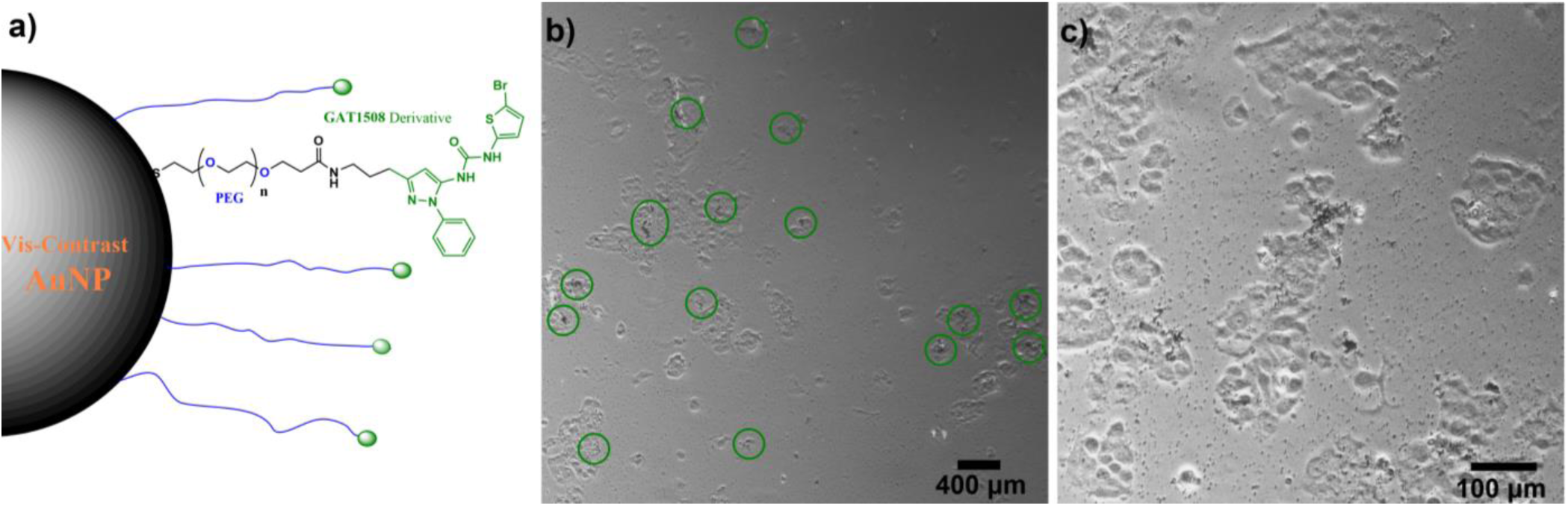
(a) Scheme of *GAT1508*-PEG-AuNPs; Optical microscope images of MCF-7 cells in the presence of *GAT1508*-coated NPs at (b) 400 nm and (c) 100 nm.

### Binding Studies of *GAT1508*-PEG-AuNPs to MCF-7 Breast Cancer Cells

To further examine potential binding of *GAT1508-*PEG-AuNPs to MCF-7 cells, an amine-terminated Alexa 594-EG_2_ derivative was synthesized (structure in **Figure S17**), since spherical AuNPs do not fluoresce. The purity of the product was confirmed by LC-MS (**Figure S17**). The fluorescent derivative was then conjugated to unreacted -COOH of *GAT1508*-PEG-AuNPs via EDC/NHS coupling, and the fluorescently-labeled NPs were purified with a Sephadex column (labeled-NPs eluted first, while unreacted Alexa 594-EG_2_ dye eluted last). ER+ MCF-7 breast cancer cells were grown at a density of 2 x 10^5^ cells/mL, and incubated with media only (control), uncoated-NPs (HOOC-PEG-AuNPs), and (Alexa 594-EG_2_)/*GAT1508-*PEG-AuNPs at 158 μg Au/mL for 20 h, followed by x2 washings with media to remove any unbound dye. Results showed absence of binding for media-treated cells (**Figure 5a,c**) and uncoated-NPs (black polygon in **Figure 5b**), and strong binding for the fluorescently-labeled *GAT1508*-NPs (**Figure 5b,d**).

**Figure 5.**
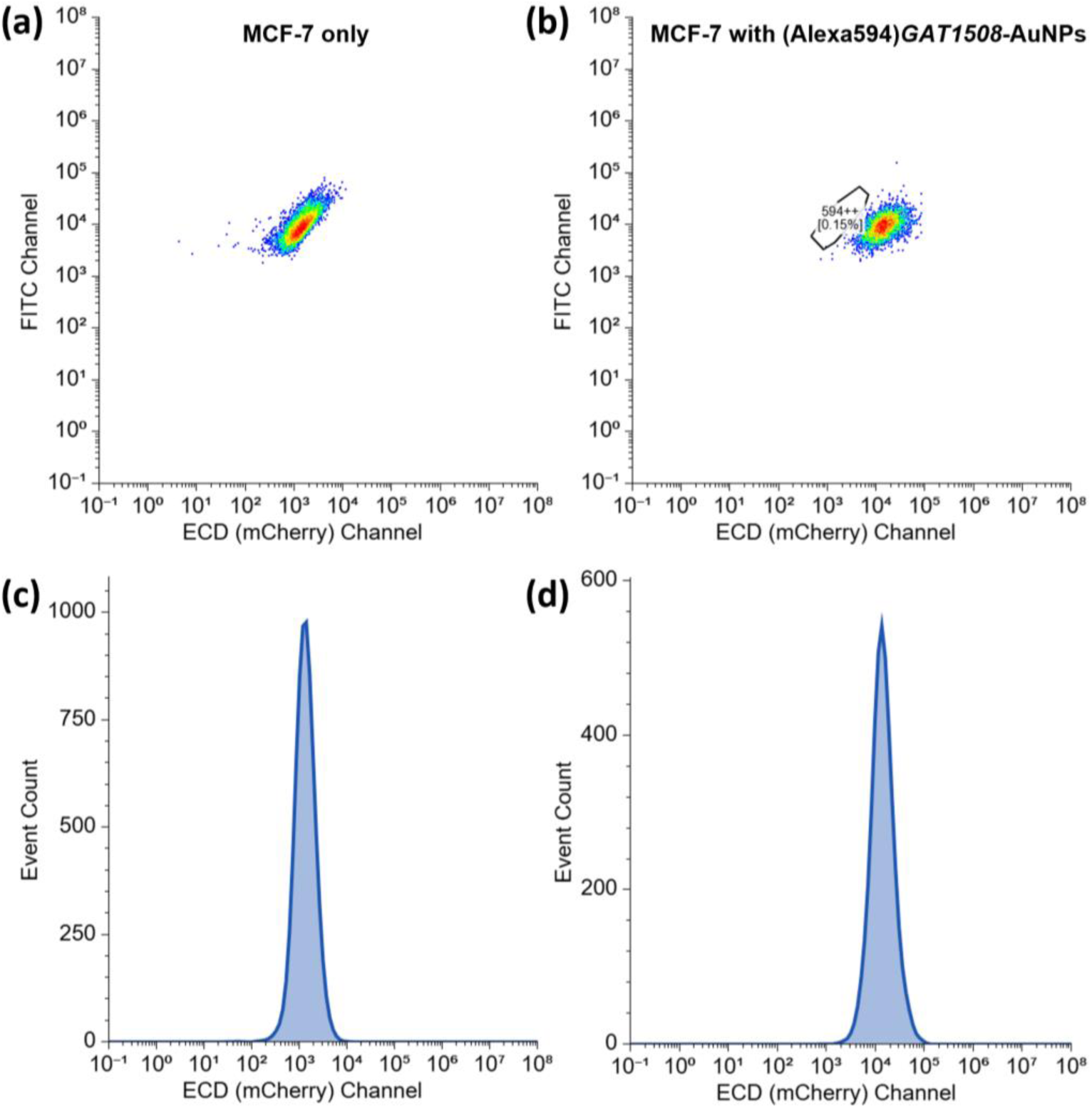
(a) Flow cytometry binding studies of MCF-7 cells incubated with (a,c) media only, and (b,d) (Alexa 594)/*GAT1508*-PEG-AuNPs at 158 μg Au/mL. A black polygon in (b) denotes absence of binding for HOOC-PEG-AuNPs to MCF-7 cells. Fluorescence patterns in the mCherry detection channel (exc./em. 587/610 nm) over the FITC channel (reference; no fluorescence) are shown in (a) and (b), while (c) and (d) represent histograms. Results are shown after removal of media and x2 washings to remove unbound dye.

The top priority in hospital breast cancer screening settings is *sensitivity* and *specificity*. Our studies indicate that our ion channel-based nanotechnology can assist in the rapid detection of ER+ breast cancer, offering selective tracing over triple (-) breast cancer cells. Most important, no fluorescent dyes were used or multiple amplification steps for detection, but a single optical microscope and a re-usable (up to 2 weeks) ion channel ligand-binding NP solution. The tracing of ER+ breast cancer through the overexpressed ion channels was feasible within hours, unlike the process of histological evaluation. Future directions involve evaluating our NP solution in tissues (histology), examining if we can differentiate malignant vs. benign tumors (e.g. lipoma) as well as ER+ from triple (-) tumors. Such low-cost and rapid technology can find wide applicability from rural areas in the US and Europe to even parts of Africa, where palpation is usually the only available source of breast cancer screening.

## Conclusions

Herein, we report the preparation of *GAT1508*-coated PEGylated AuNPs as an optical microscope bio-marker for ER+ breast cancer screening, taking advantage of the overexpressed GIRK1 channels in ER+ human breast cancer cells. Two derivatives of GAT1508 (urea-based molecule) were synthesized and characterized in detail. However, only GAT1508-PA (alkylamine chain extension from the pyrazole ring) showed partial inhibition of GIRK1/2-mediated K^+^ currents in transfected HEK293^GIRK1^ cells using three characterization methods: TEVC, MPC (whole-cell), and the TFA. Computational docking studies predicted strong binding for the GAT1508 propylamine derivative, both in amine form (GAT1508-PA) and in amide form (GAT1508-PA-EG2; the ligand is conjugated to the NP via a PEG linker). A *GAT1508*-NP of ∼4 nm was then synthesized. UV-Vis studies revealed the presence of the conjugated ligand at 260 nm in addition to the SPR of AuNPs at 526 nm. Flow cytometry studies showed binding of Alexa 594-labeled *GAT1508*-PEG-AuNPs in ER+ MCF-7 breast cancer cells and strong interaction. Finally, incubation of fixed MCF-7 cells with a *GAT1508*-NP solution, followed by several washings to remove unbound NP, led to detection of breast cancer cells using a simple optical microscope, without the need for fluorescent dyes and/or additional amplification steps. Detection was not feasible in triple (-) breast cell line MDA-MB-231 that does not express GIRK1. This is the first study, to our knowledge, that highlights the importance of nanotechnology as a tool of detection of overexpressed ion channels in breast cancer. Future direction will involve evaluating the potential of this nanoprobe in detecting cancerous mammary gland tissues (histology) and potentially differentiating malignant vs. benign tumors in excised organs of ER+ orthotopic xenograft rodent models.

## Materials and Methods

### Materials

5-bromothiophene-2-carboxylic acid, diphenylphosphorylazide (DPPA), and triethylamine were purchased from TCI chemicals. 3-(3-aminopropyl)-1-phenyl-1H-pyrazol-5-amine*HCl was purchased from Santa Cruz Biotechnology. 1-(3-aminomethyl)phenyl-3-methyl-1H-pyrazol-5-amine was purchased from Ambeed. GAT1508-PA and GAT1508-PA-EG2 were synthesized in house. Dodecanethiol-functionalized gold nanoparticles and carboxymethyl-PEG-thiol (HOOC-PEG-SH; 5,000 g/mol) were purchased from Nanoprobes Inc and Laysan Bio Inc. respectively. *GAT1508*-PEG-AuNPs were synthesized in house. HEK-293T epithelial-like cells (CRL-3216^™^), MCF-7 human breast cancer cells (mammary gland; HTB-22), MDA-MB-231 breast cancer cells (mammary gland; HTB-26), and DMEM media were acquired from ATCC. Fetal bovine serum (FBS) was purchased from ThermoFisher. Alexa Fluor™ 594 NHS (succinimidyl ester) was purchased from Lumiprobe. Alexa 594-EG_2_-amine was synthesized in house.

### Synthesis of Urea Ligands GAT1508-EA and GAT1508-PA

A 2 mL anhydrous toluene suspension of 5-bromothiophene-2-carboxylic acid (2 eq.) and alkylated amine of benzyl pyrazole-amine (in salt form; 1 eq.) were placed in a microwave vial along with triethylamine(3 eq.) and DPPA (1.5 eq.) and irradiated for 5 mins at 120 °C. The reaction mixture was poured into water (pH=6) and extracted with ethyl acetate. The combined organic layers were washed with water and brine, dried over Na_2_SO_4_ and evaporated under vacuum. The crude residue was purified by preparative HPLC to isolate the urea.

*GAT108-EA*: ^1^H NMR, (400 MHz, Methanol-D4, Reference = δ 3.31): δ 2.15 (3H), δ 4.44 (2H), δ 5.42 (1H), δ 6.25 (1H), δ 6.77 (1H), δ 7.33 (1H), δ 7.40 (1H), δ 7.46-7.47 (2H). MS (ESI): *m/z*: 405 ([M+H]^+^). *GAT108-PA*: ^1^H NMR, (400 MHz, Dimethyl Sulfoxide-D6, Reference = δ 2.50): δ 1.71 (2H), δ 2.45 (2H), δ 3.12 (2H), δ 5.25 (2H), δ 6.20 (1H), δ 6.40 (1H), δ 6.83 (1H), δ 7.26 (1H), δ 7.44 (2H), δ 7.56 (2H), δ 9.67 (1H). MS (ESI): *m/z*: 420 ([M+H]^+^).

### Synthesis of HOOC-PEG-Gold Nanoparticles (HOOC-PEG-AuNPs)

PEGylated AuNPs were prepared by ligand exchange of C12-S-AuNPs with excess HOOC-PEG-SH. 1000 mg of HOOC-PEG-SH (0.2 mmol, *M*_n_ = 5,000 g/mol per manufacturer) were dissolved in 30 mL DCM and equilibrated at RT. Subsequently, a 20 mL DCM solution of C12-S-AuNPs (320 mg, 89% metal per TGA = 285 mg Au) was added dropwise to HOOC-PEG-SH (the polymer concentration was reduced to 2 wt%), followed by continuous stirring at RT overnight. The solvent was then evaporated; the mixture was dispersed in the minimum amount of DCM (∼2 mL) and precipitated in hexane to remove liberated dodecanethiol. The crude product was then dispersed in DI water at 10 wt% and centrifuged at 20,000 rpm for 3 h at 8 °C to remove unbound PEG-SH (supernatant), concentrating the HOOC-PEG-AuNP precipitate. The final pellet (∼278 mg) was collected and dried under vacuum overnight at RT. Every step was monitored by TGA. The collected PEG-coated AuNPs were highly soluble in water. Addition of toluene showed clear transfer to the aqueous phase, unlike C12-S-AuNPs.

### Preparation of *GAT1508-*PEG-AuNPs

HOOC-PEG-S-AuNPs (50 mg, 35 wt% PEG, => 17.5 mg PEG in Au, => 3.5 ×10^-6^ moles –COOH since PEG is 5K) were dispersed in 1 mL DMF (4 wt%) and sonicated for 10 mins. Subsequently, EDC (2.2 mg, 14 ×10^-6^ moles, 155.25 mg/mmol, **4 eq**.) was added and was left to react under stirring. After 10 mins, NHS (1.6 mg, 14 ×10^-6^ moles, 115.09 mg/mmol, **4 eq**.) was added. The mixture was sonicated for 5 mins and was left to react for 20 mins. After, GAT1508-PA (**5.9 mg**, 14 ×10^-6^ moles, 419 mg/mmol, **4 eq.**) was added in 1.5 mL DMSO and the reaction mixture was left to react overnight. It was then dialyzed over DI water for 3-4 days (replacing twice per day the water) to remove free drug and lyophilized.

### Hydrolysis of *GAT1508*-PEG-AuNPs / Ligand Quantification

2 mg of *GAT1508*-PEG-AuNPs, 2 mg of HOOC-PEG-AuNPs, and 1 mg of GAT1508 were dispersed in 6 N HCl and heated overnight at 100 °C to break the amide bonds and release the ligand. After, the solvent was evaporated, and samples were dissolved in 1 mL DMSO. In the case of NPs, a centrifugation step at 14,000 rpm was performed to separate the NPs (pellet) from the supernatant which contained the free ligand. Quantification was performed based on calibration curve of the ligand peak at 261 nm using UV-Vis (Nanodrop). The absorbance of the supernatant of GAT1508-NPs was subtracted from the one of uncoated-NPs.

### Synthesis of Alexa Fluor 594-(Ethoxy)Ethanamine (Alexa Fluor 594-EG_2_-NH_2_)

1,2-Bis(2-aminoethoxy)ethane (0.18 mg, 1.21 x 10^-6^ moles, MW= 148.2 g/mol, 3 eq.) was dissolved in 0.2 mL dry DMSO. Then Alexa Fluor 594 NHS ester (0.33 mg, 2 mM, 0.4 x 10^-6^ moles, 1 eq, MW = 819.9 g/mol) was dissolved in 0.2 mL dry DMSO and added dropwise. The reaction was left overnight (covered with foil). The product was purified by precipitation in large volume of diethyl ether to dissolve excess diamine, and characterized by ESI-MS spectroscopy. Initial reactants were not observed by ESI-MS. ESI-MS m/z calculated for C_42_H_51_N_4_O_11_S_2_: 851.30 (M+H)^+^; Found: 851.0 (M+H)^+^.

### Conjugation of *GAT1508-*PEG-AuNPs with Alexa Fluor 594-EG_2_-Amine (Synthesis of Alexa 594/*GAT1508*-coated NPs)

*GAT1508-*NP (11 mg; 39 wt% PEG, => 4.3 mg PEG in Au, => 0.86 ×10^-3^ mmoles-COOH since PEG is 5K; assuming all -COOH are reactive; 1 eq.) was dispersed in 627 μL dry DMSO and sonicated until well-dispersed. Subsequently, 23 μL of a 26 mg/mL solution of EDC*HCl (0.59 mg, 3.1×10^-3^ mmoles, 191.70 g/mol, 4 eq., 2.6 mg in 0.1 mL) in dry DMSO were added. After 5 mins, 17 μL of a 21 mg/mL NHS solution (0.36 mg, 3.1×10^-3^ mmoles, 115.09 g/mol, 4 eq., 2.1 mg in 0.1 mL) in dry DMSO were added. The mixture was sonicated for 3 mins and was left to react for 20 mins. The dispersion was added dropwise to a 433 μL solution of Alexa Fluor 594-EG_2_-amine (0.33 mg, 0.39×10^-3^ mmoles, 850.29 g/mol, 0.5 eq.) in dry DMSO, and the reaction was left to react overnight (final C_nanomaterial_ ∼ 1 wt%, 1100 μL). The unreacted dye was separated from the labeled-AuNPs using a Sephadex G-25 column, where free dye could be detected in a separate fraction. The material purity was tested with TLC to confirm absence of unreacted dye.

### *X. laevis* Oocyte Expression

Plasmid DNAs of GIRK channel subunits were linearized prior to *in vitro* transcription. Capped RNAs were transcribed using mMESSAGE mMACHINE T7 transcription kit (Thermo Fisher Scientific). *Xenopus* oocytes were surgically extracted, dissociated, and defolliculated by collagenase treatment and microinjected with 50 nL of a water solution containing the relative amount (in ng) of each GIRK subunit RNA. For TEVC experiments, oocytes were kept 2 days at 17 °C before recording.

### Two-Electrode Voltage-Clamp Experiment (TEVC)

Whole-oocyte currents were measured by TEVC with GeneClamp 500 (Molecular Devices) or TEC-03X (NPI) amplifiers. Electrodes were pulled using a Flaming-Brown micropipette puller (Sutter Instruments) and were filled with 3 m KCl in 1.5% (w/v) agarose to give resistances between 0.5 and 1.0 MΩ. The oocytes were bathed in ND96 recording solution comprising: KCl 2 mM, NaCl 96 mM, MgCl_2_ 1 mM, and HEPES 5 mM, buffered to pH 7.4 with KOH. Where indicated, GIRK channel currents were assessed in a high-K^+^ recording solution comprising (in mm): KCl 96 mM, NaCl 2 mM, MgCl_2_ 1 mM, and HEPES 5 mM, buffered to pH 7.4 with KOH. Currents were digitized using a USB interface (National Instruments) and recorded using WinWCP software (University of Strathclyde). To study GIRK channels, oocytes were held at 0 mV, and currents were assessed by 100-ms ramps from −80 to +80 mV that were repeated every second. The effect of different ligands was determined at −80 mV and then the channels were blocked by 5 mm BaCl_2_. Block was expressed as the percent-current block normalized to the maximum current. Between 8 and 12 oocytes from different *Xenopus* frogs were studied per experiment.

### Culture of HEK293 Cells

HEK293-T cells were obtained from the ATCC and maintained in Dulbecco’s modified Eagle’s medium supplemented with 10% fetal bovine serum and 1% penicillin and streptomycin (HyClone). For patch-clamp studies, cells were seeded on glass coverslips and transfected 24 h later using a polyethyleneimine solution (1 mg/mL) at a ratio of 8 μL per μg of DNA. To study GIRK currents, cells were transfected with 0.75 μg each of plasmids encoding GIRK1 and GIRK2.

### Manual (Whole-Cell) Patch Clamp (MPC)

Whole-cell currents were recorded with an Axopatch^TM^ 200B amplifier (Molecular Devices) controlled via a USB-interface (National Instruments) using WinWCP software (University of Strathclyde). Currents were acquired through a low-pass Bessel filter at 2 kHz and were digitized at 10 kHz. Patch pipettes were fabricated from borosilicate glass (Clark), using a vertical puller (Narishige) and had a resistance of 2.5–4 MΩ when filled with an intracellular buffer comprising: 140 mM KCl, 2 mM MgCl_2_, 1 mM EGTA, 5 mM Na_2_ATP, 0.1 mM Na_2_GTP, and 5 mM HEPES, pH 7.2. Cells for study were selected based on GFP expression using an epifluorescence microscope (Nikon). To study the activity of GIRK channels, cells were held at 0 mV, and currents were assessed by ramps from −80 to +80 mV that were repeated at 1 Hz. Cells were perfused via a multichannel gravity-driven perfusion manifold with a physiological buffer comprising: 135 mM NaCl, 5 mM KCl, 1.2 mM MgCl_2_, 1.5 mM CaCl_2_, 8 mM glucose, and 10 mM HEPES, pH 7.4, and then quickly transitioning to a high-K^+^ buffer comprising 5 mM NaCl, 135 mM KCl, 1.2 mM MgCl_2_, 1.5 mM CaCl_2_, 8 mM glucose, and 10 mM HEPES, pH 7.4. The barium-sensitive component of the current, observed when cells were perfused with the high-K^+^ buffer, was analyzed and determined by perfusing 0.1 mM BaCl_2_ in the high-K^+^ buffer at the end of each experiment. HEK293 cells had a mean whole-cell capacitance of 10 ± 1 picofarads; series resistance was typically <10 MΩ, and the voltage-error of <3 mV was not adjusted.

### Thallium Flux Assay (TFA)

HEK-293T cells were used for transient expression of human GIRK1 and GIRK2 channels. Cells were maintained in DMEM (ATCC, VA) supplemented with 10% (v/v) fetal bovine serum (Thermo Fisher Scientific, Waltham, MA) and 0.5% (v/v) Penicillin-Streptomycin (Gibco, NY) at 37 °C in a humidified incubator with 5% CO₂. Cells were passaged the day before electroporation to reach late-log phase on the day of the experiment. For transient transfection, HEK-293T cells were electroporated with pXOOM plasmids encoding human GIRK1 and GIRK2 (Origene, Rockville, MD). Electroporation was performed in 0.2 cm cuvettes using a Bio-Rad GenePulser MXcell^TM^ system with Cytomix buffer composed of (final concentrations): 120 mM KCl, 0.15 mM CaCl₂, 10 mM K₂HPO₄/KOH (pH 7.6), 2 mM EGTA, 25 mM HEPES/KOH (pH 7.6), 5 mM MgCl₂, and freshly supplemented on the day of experiment with 2 mM ATP and 5 mM glutathione.^43^ All solutions were adjusted to pH 7.6 with KOH, filtered through a 0.22 μm filter, and ATP/glutathione were added immediately before use to prevent degradation. Following electroporation, cells were recovered in complete culture medium and plated into poly-D-lysine–coated, black-walled, clear-bottom 96-well plates (Corning) at 80,000 cells/well in 100 μL/well and incubated overnight at 37 °C, 5% CO₂ to allow channel expression.

On the next day, FLIPR Potassium Assay Kit (Molecular Devices, Cat#. R8222) was used to measure GIRK channel activity. Cell plates were removed from the incubator and loaded with an equal volume (100 uL/well) of Loading Buffer (prepared per kit instructions) consisting of acetoxymethyl (AM) ester tagged Tl+ indicator dye, and masking dye dissolved in 1x Hank’s Balanced Salt Solution (HBSS) pH 7.4, and 20 mM HEPES. Plates were incubated for 1 h at room temperature in the dark to facilitate dye loading. To introduce Tl+/ligand into the cell plate, 5x solution consisting of different concentrations of GAT1508/PA and 3 mM Tl+ in chloride-free buffer was prepared and aliquoted in 96-well v-bottom microplates (Greiner, 651201).

Cell microplates and compound plates were then transferred to the Flexstation 3 instrument (Molecular devices, CA). Baseline fluorescence was acquired for 10 s with excitation of 485 ± 20 nm, emission of 525 ± 30 nm, PMT sensitivity of 6, and an addition rate of 3. After the baseline recording, 50 μL/well of compound solution was added automatically by the FlexStation pipetting head at 30 s timepoint, and recording continued for a total of 5 min. Test compounds included GAT1508 (0.03 μM–10 μM) and GAT1508-PA (10 μM, 30 μM). Blank samples are electroporated cells which received the same buffer with (0.05% DMSO final concentration) in place of GAT compounds. Fluorescence data were normalized to baseline (ΔF/F) for each well. For the analysis shown in **Fig. 1f**, the percentage change in fluorescence (ΔF/F %) was quantified at 30 s after compound addition. Each condition was tested in triplicate across at least three independent experiments. Data are presented as mean ± SEM (Standard Error of the Mean).

### Molecular Docking

GAT1508-PA and GAT1508-PA-EG2 were docked into the binding pocket of GIRK1/2 channel using induced-fit-docking (IFD) simulations (Maestro, version 2025-3). For IFD simulations, default parameters were used. For redocking and ranking of the docked poses, XP scores were used. The pose of the docked ligand was selected by comparing it to GAT1508 parent compound which was consistent with cryo-EM density for the GAT1508 binding site.

### Binding Free Energy Calculations

The Prime MM-GBSA module of Schrodinger’s Maestro was used to calculate the binding free energy of GAT1508-PA and GAT1508-PA-EG2 in complex with GIRK1/2 channel. An implicit solvation model VSGB 2.0 is implemented in this module to calculate the optimized energy of free ligand, free receptor, and a receptor-ligand complex. The binding free energy (dG_bind) was calculated using Formula:

> dG_bind = Energy of the complex – (Energy of ligand + Energy of protein)

where dG_bind represents the approximated binding free energy. It is the estimated energy difference between the bound state and sum of energies of unbound states.

### Gating Molecular Dynamics (GMD) Simulations

Amber24 was used to conduct simulations, applying the Amber ff19SB and Lipid21 force fields. Ligand geometries and charges were generated at the B3LYP/CC-PVTZ level using Gaussian 16. CHARMM-GUI server (http://www.charmm-gui.org)^44^ was used for membrane building and generating the final system with ions and water. The GIRK1/2-PIP2 and ligand were immersed in the lipid bilayer of POPC, POPS, POPE and cholesterol at a ratio of 25:5:5:1, respectively.^28^ TIP3 water model was used to solvate the system. KCl was used to neutralize the system at 150 mM. System was minimized in two steps. To bring the system to 300K, heating was performed in 3 steps. In the final phase, the system was equilibrated and relaxed. The production phases for these simulations consisted of 500 ns with a 3fs time step. Simulations were run in triplicate on the systems containing a compound. The results of the GMD simulations were analyzed using various tools and methods, including the built-in utility of the GROMACS program from Groningen University, as well as in-house scripts.

### Optical Microscope Breast Cancer Cells Binding Studies with *GAT1508*-PEG-AuNPs

ER+ MCF-7 breast cancer cells and triple (-) MDA-MB-231 cells were cultured in DMEM medium supplemented with 10% FBS in a 25 cm^2^ *tissue-culture treated flask* for 5-6 days and were then trypsinized. 1 mL of that culture was transferred (at a density of 2 x 10^5^ cells/mL) to three 12-well plates, followed by 1 mL of media. After two days of subculturing, media was decanted, and 2 mL of fresh media (control), or HOOC-PEG-AuNPs (uncoated-NPs) in media, or *GAT1508*-PEG-AuNPs in 2 mL media at 158 μg Au/mL were added. After 20 h of incubation, media (with and without NPs) was decanted to remove unbound NPs, cells were washed x2 with media to remove unbound NPs, and imaged under an optical microscope. A slightly different procedure involved culturing MCF-7 cells in adherent glass *coverslips* at a density of 2-5 x 10^5^ cells/mL, followed by fixing with 4% formaldehyde, washing, and incubating the cells in a *GAT1508*-NP solution for 10-18 h, followed by washing steps to remove unbound NPs, and optical imaging. The latter procedure enabled the preservation of the MCF-7 cell structure, since these cells are known to detach from the adherent surface or alter their shape in the presence of PBS or shear stress.^45^ At least 10 mins incubation period per washing step was used to remove unbound NPs.

### Breast Cancer Cells Binding Studies with (Alexa 594)/*GAT1508*-PEG-AuNPs

ER+ MCF-7 breast cancer cells were first cultured in DMEM medium supplemented with 10% FBS in a 25 cm^2^ tissue-culture treated flask for 5-6 days and were then trypsinized. 1 mL of that culture was transferred (at a density of 2.0 x 10^5^ cells/mL) to three 12-well plates, followed by 1 mL of media. After two days of subculturing, media was decanted, and 2 mL of fresh media (control), or HOOC-PEG-AuNPs in media, or (Alexa 594)/*GAT1508*-PEG-AuNPs in 2 mL media at 158 μg Au/mL were added. After 20 h of incubation, media (without and with NPs) was decanted to remove unbound NPs, cells were washed with media, and trypsinized. Cells were then resuspended in 1 mL fresh DMEM media, mixed well to break any cell clusters, and passed through a 100 μm Falcon® cell strainer. Flow cytometry studies were then conducted using only cell singlets and monitoring the same polygon area of interest (gating) for consistency in all examined samples. Fluorescence was measured on the mCherry channel (ECD; exc./em. 587/610 nm) over the FITC channel (exc./em. 495/519 nm) as a reference. Alexa 594 has an excitation/emission at 590/617 nm.

### Characterization

A Biotage Initiator **microwave system** was used for the synthesis using a microwave vial. Reaction progress was monitored by thin-layer chromatography (TLC). **NMR** was acquired in a 500 MHz Bruker Avance Neo using d-MeOH as the solvent. **Preparative HPLC** was used to purify the compounds. Those were dissolved in DMF and injected into a Phenomenex Luna 5 µm C18(2) 100 Å LC Column, 150 x 21.2 mm, AXIA Packed (Part no: 00F-4252-P0-AX) at a flow rate of 20 mL/min. Solvent A was water in 0.1 M formic acid and Solvent B was acetonitrile in 0.1 M formic acid. For **LC-MS**, samples were analyzed on an Agilent ZORBAX RRHT StableBond-C18 column (1.8 μm, 2.1 × 50 mm, part no: 827700-902) at a flow rate of 0.600 mL/min with the following HPLC method: 0 min 90/10 A/B, 3.5 min 10/90 A/B, 5.5 min 10/90 A/B, 6.0 min 90/10 A/B wherein A is water (+0.1 % formic acid) and B is acetonitrile (+0.1 % formic acid). All final compounds are >95 % pure by HPLC analysis. For **TGA** analysis, the samples were vacuum-dried overnight at room temperature before the analysis and measured (in triplicate) on a Mettler Toledo TGA thermal analyzer at a heating rate of 20 °C/min under N_2_, from 50–600 ^◦^C. **UV-Vis** images were collected on a DeNovix DS11-FX Nanodrop between 220-750 nm. **TEM** images were collected on a Philips CM12 electron microscope operated with an acceleration voltage of 120 kV. C12-S-AuNPs were prepared at 0.5 mg/mL in hexane, while *GAT1508*-PEG-AuNPs were prepared in water at 0.5 mg/mL. To estimate core size, more than 300 NPs were measured and analyzed with ImageJ. **Optical microscopy** studies were collected on an inverted microscope. **Flow cytometry** was conducted in a Beckman Coulter CytoFLEX S cell analyzer (CILS Core Facility, Northeastern University). Analysis was performed using Floreada software. For that, the FLIPR potassium assay kit from Molecular Devices was used.

## SUPPLEMENTARY MATERIALS

Figures **S1-S17** are available online.

## ACKNOWLEDGMENTS

This research was funded by NIH R01HL059949-28 and NIH R01NS131467-03. We would like to thank Prof. Roman Manetsch (NU) for allowing us to use his preparative HPLC and LC-MS. Dr. Guoxin Rong (CILS Core Facility, Northeastern University) is also acknowledged.

