## Supplementary Figures S1-S17 for "Ion Channel Nano-Diagnostics for ER+ Breast Cancer"

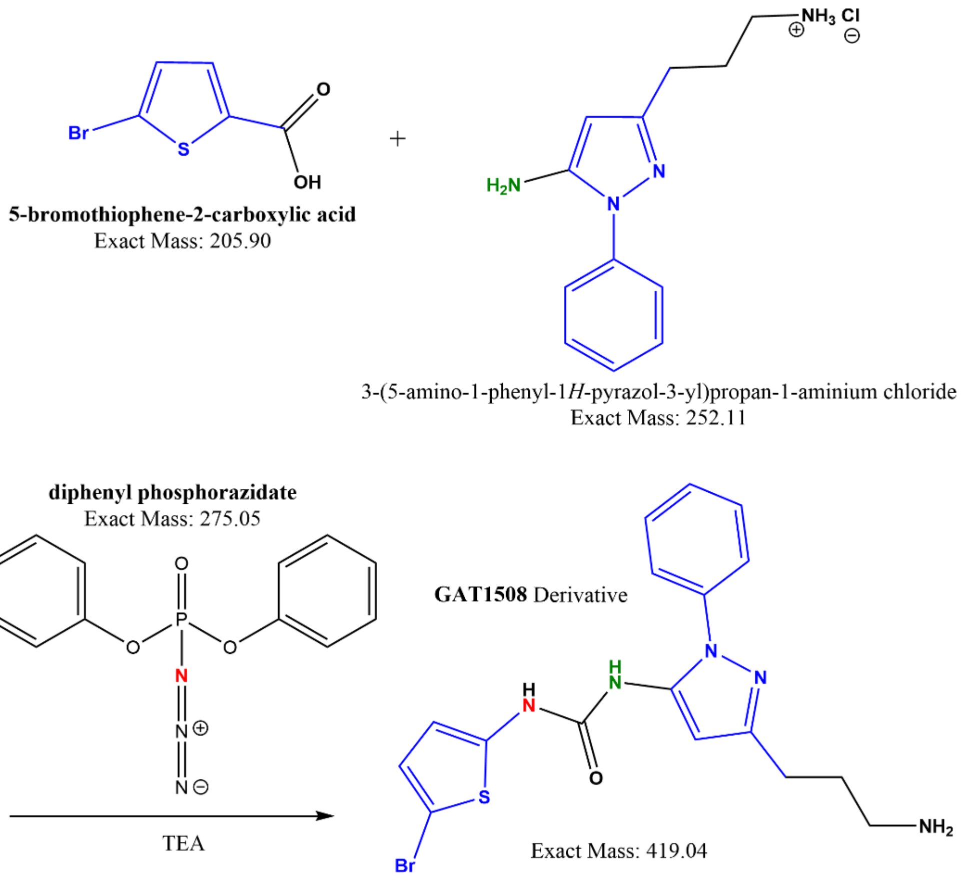


**Scheme S1**. Reaction between 5-bromothiophene-2-carboxylic acid and benzyl pyrazole alkylated amines in the presence of TEA and DPPA.

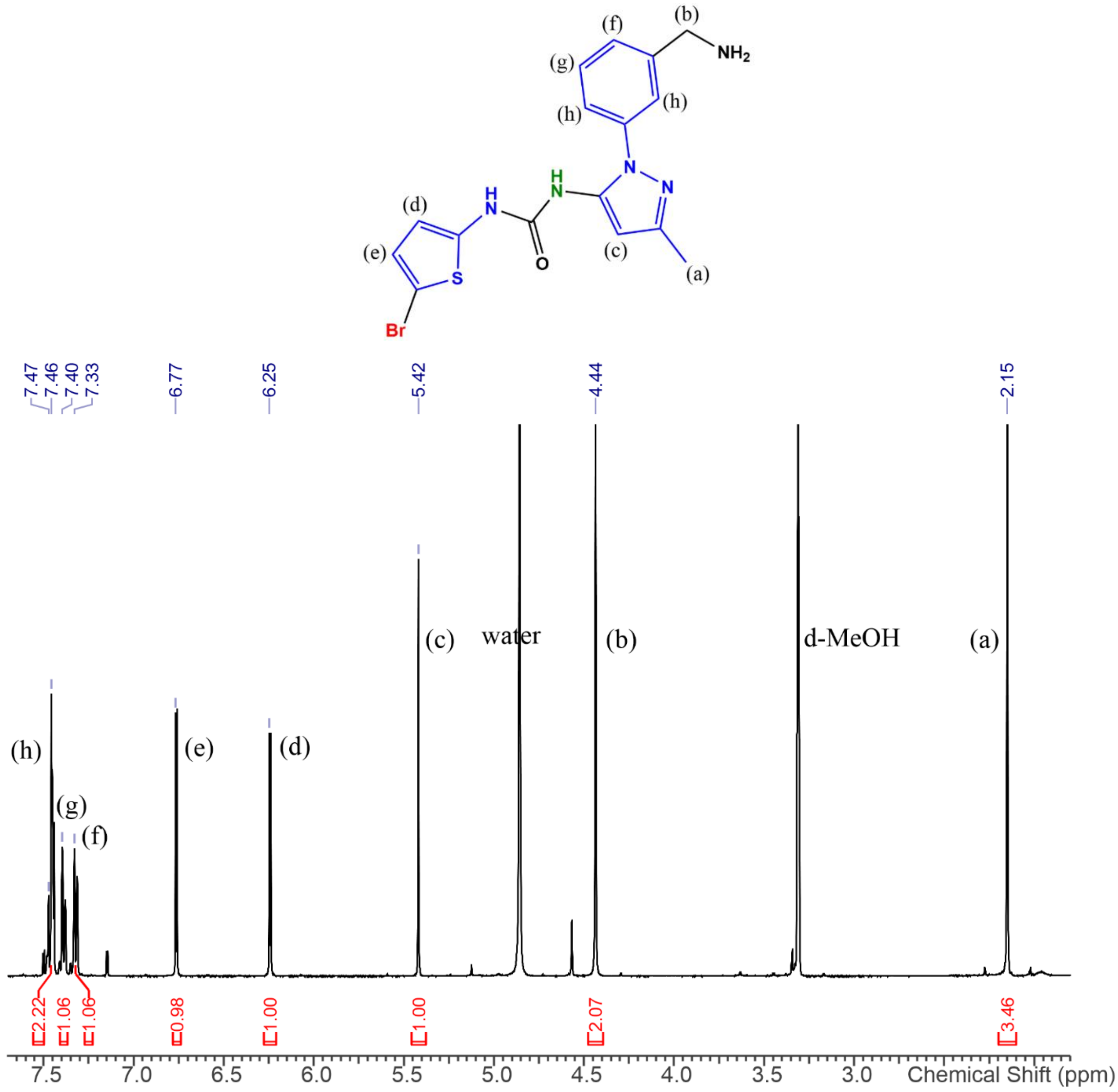


**Figure S1**. NMR of GAT1508 ethylamine in d-MeOH. The urea protons rapidly exchange with d-MeOH and thus cannot be seen.

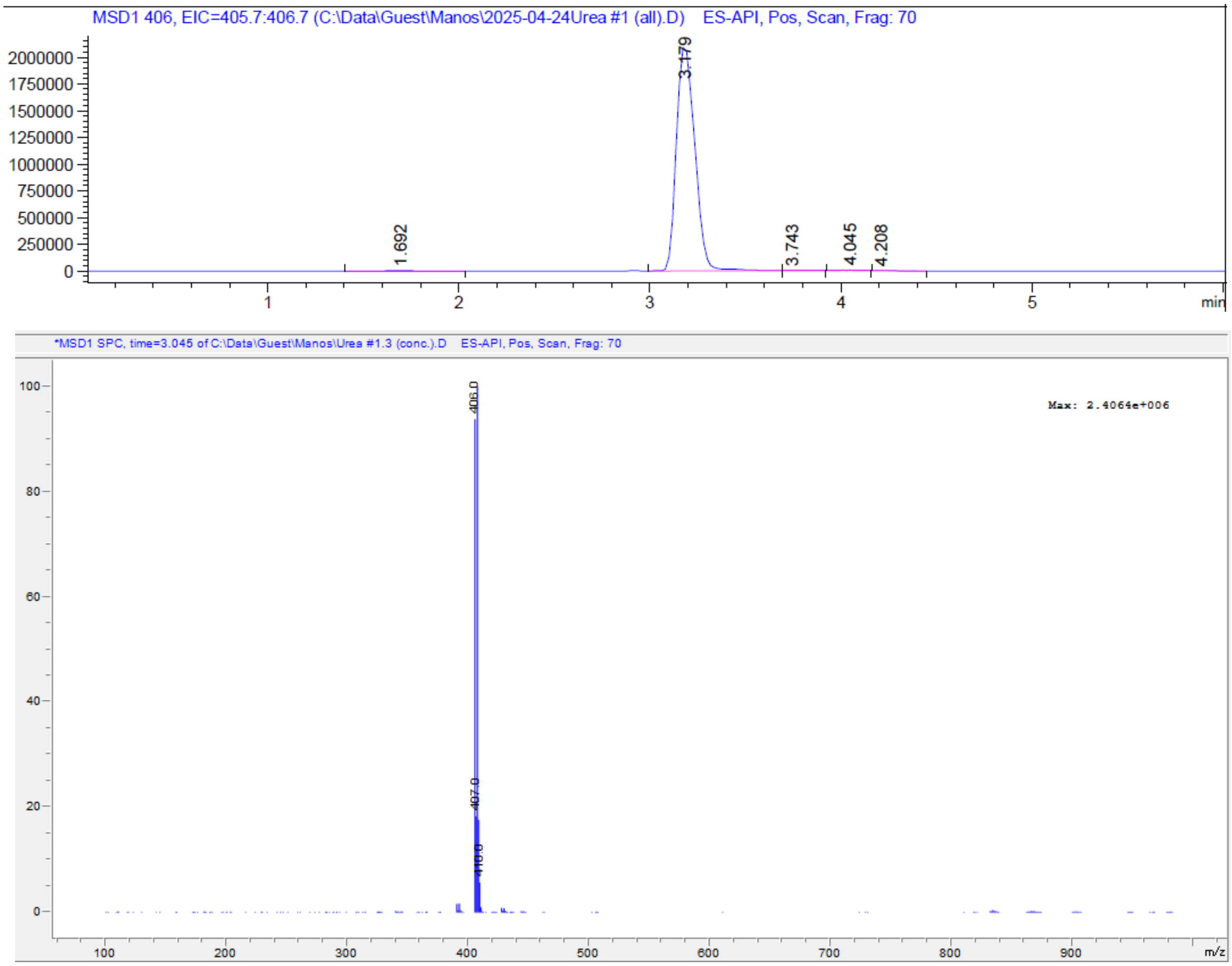


**Figure S2**. MS of GAT1508 ethylamine with a molecular weight of 405 g/mol.

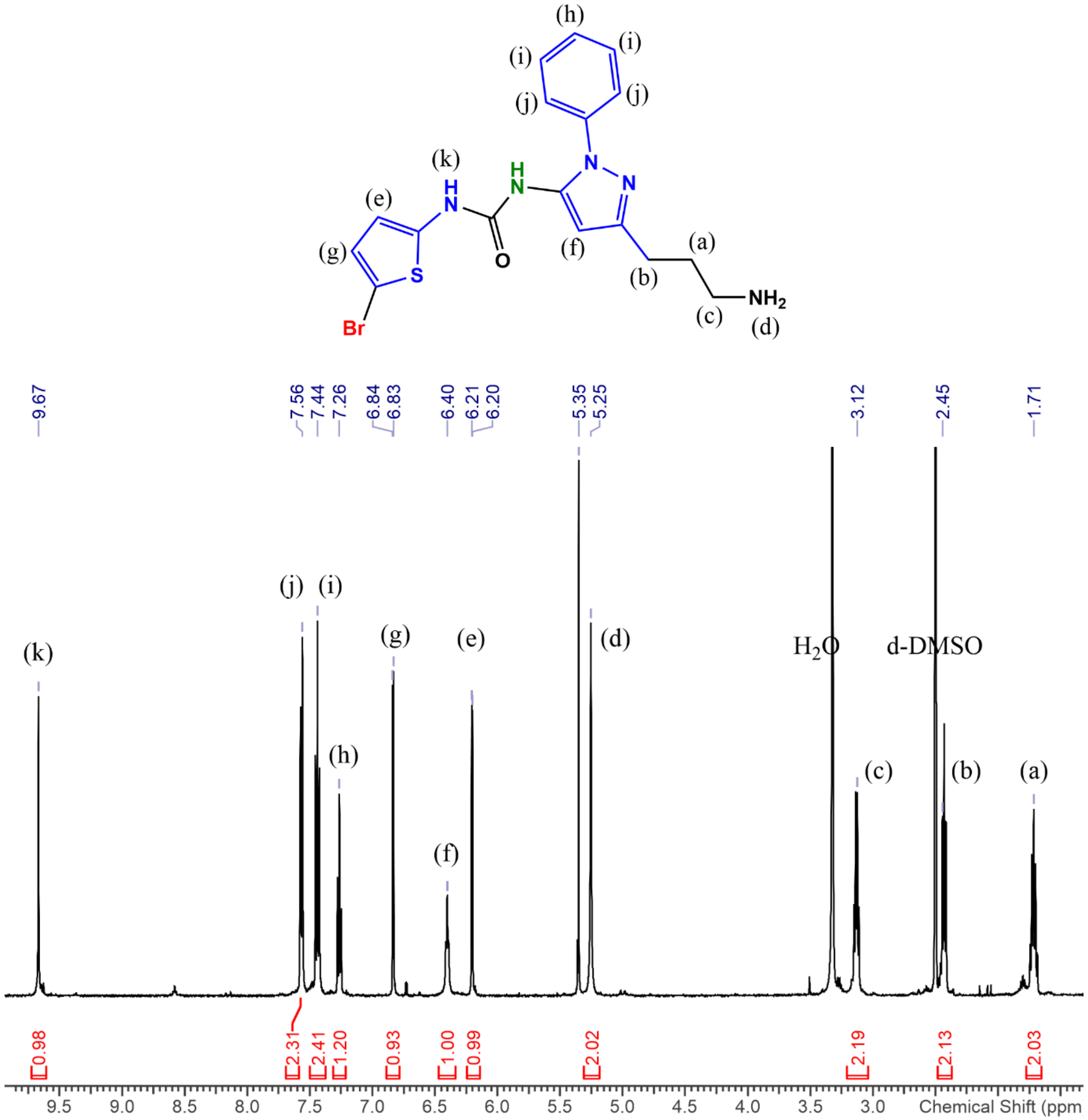


**Figure S3**. NMR of GAT1508 propylamine in d-DMSO. Only one of the urea protons can be seen. This is reported in the literature with high hydrophilic compounds.

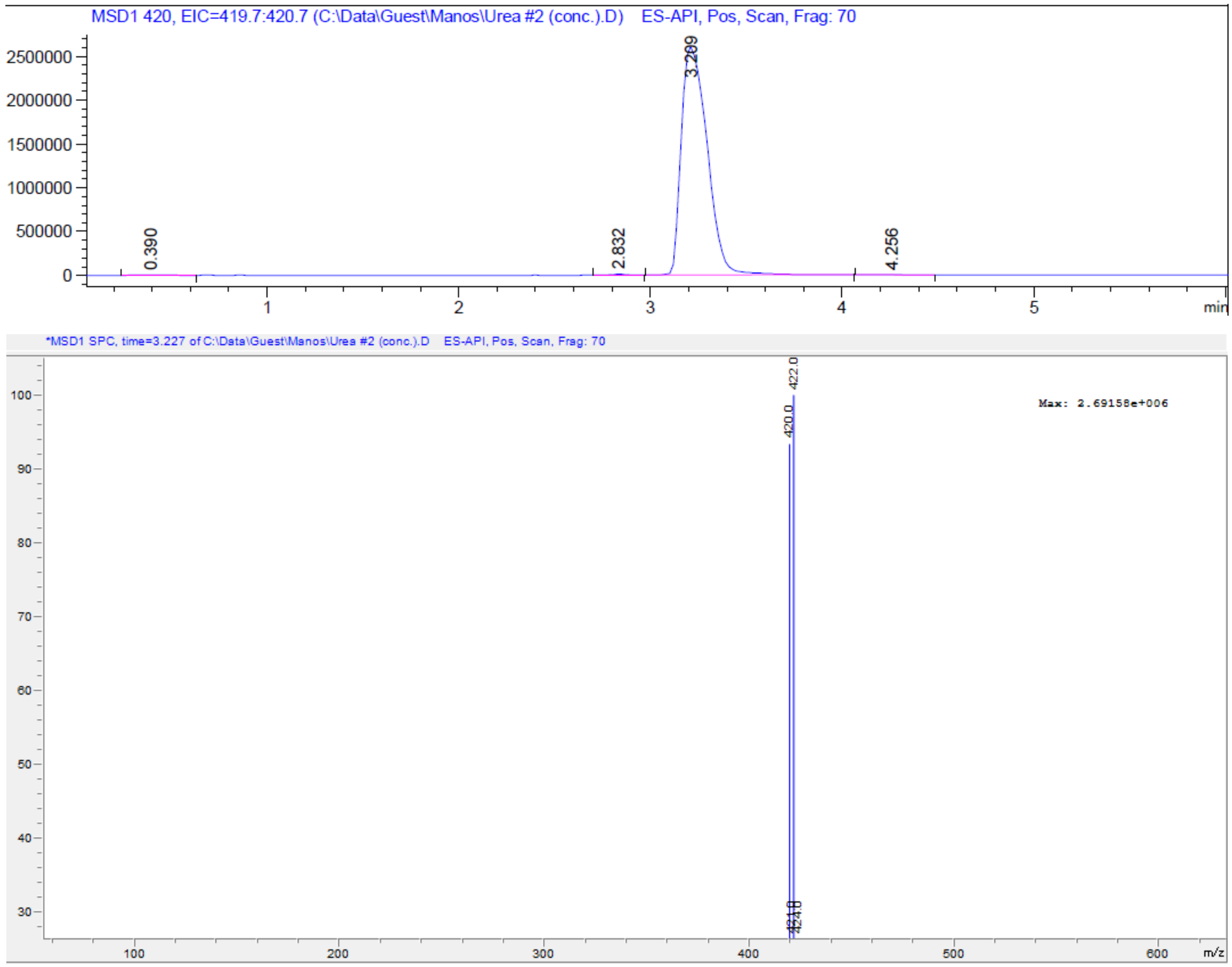


**Figure S4**. MS of purified GAT1508 propylamine with a molecular weight 419 g/mol.

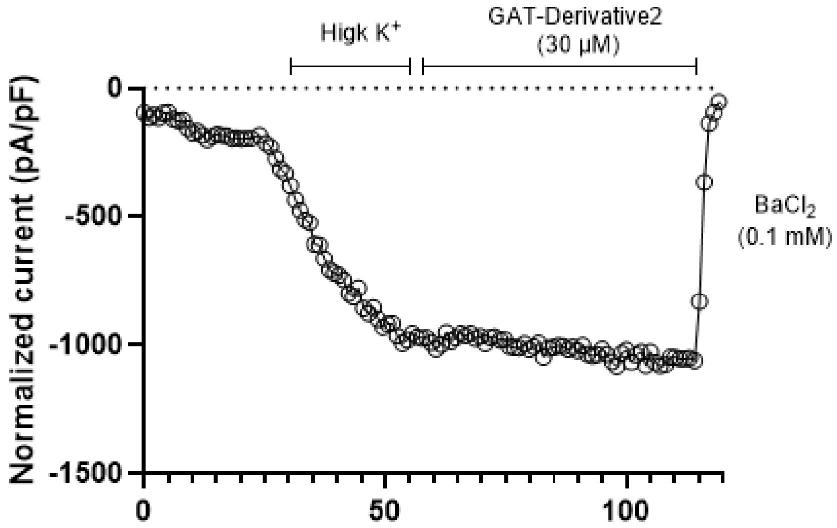


**Figure S5**. Whole-cell patch clump of the GAT1508 ethylamine showing absence of interaction with the GIRK1/2 channels overexpressed in HEK293 cells.

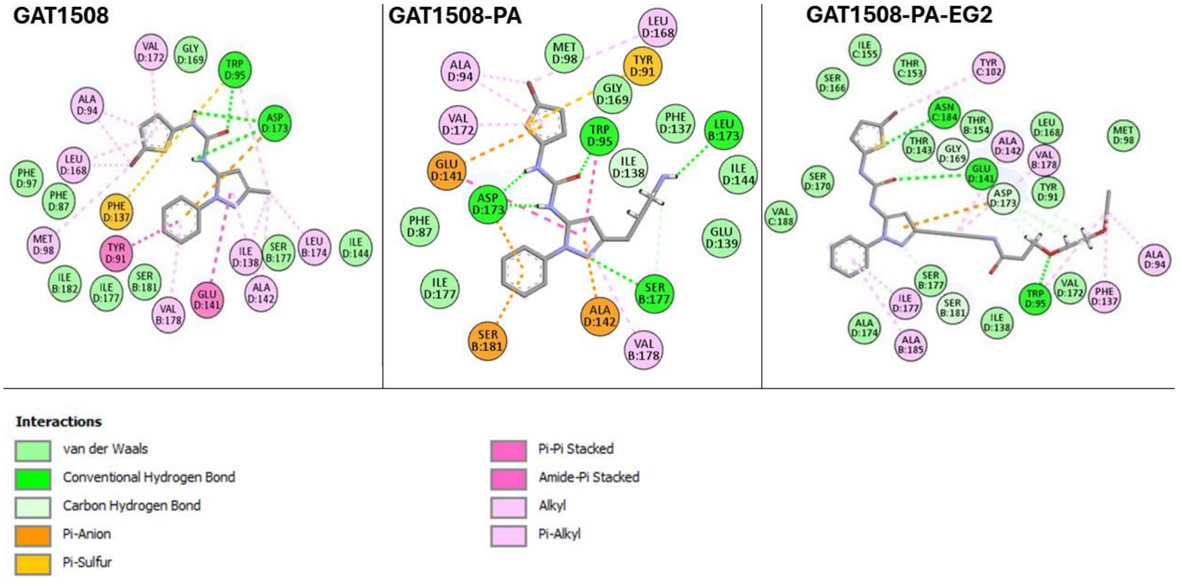


**Figure S6**. 2D interaction diagram of GAT1508 and GAT1508-EG2 showing that Asp-173 and Trp-95 are the crucial residue interactions for keeping the ligands stable in the binding pocket.

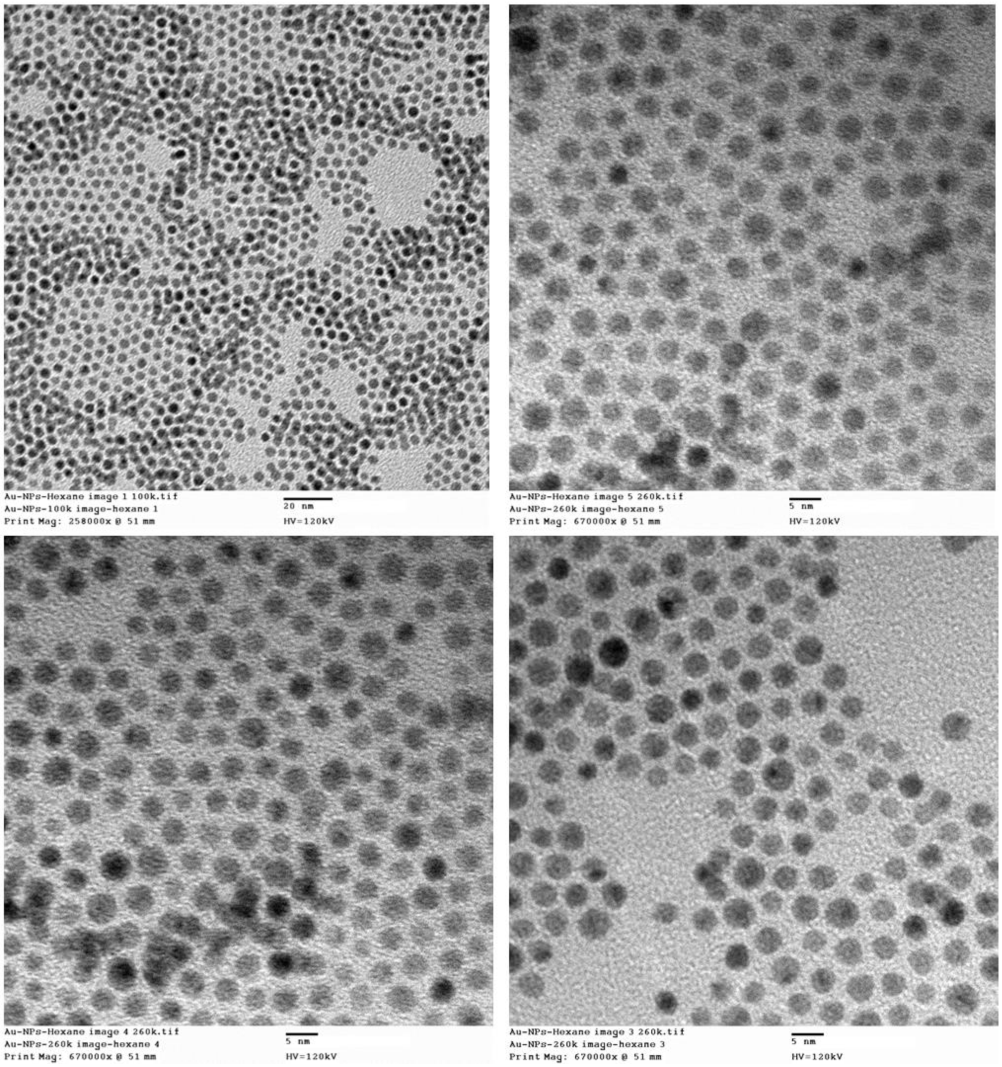


**Figure S7.** TEM of ~3.7 nm **dodecathiol-coated AuNPs** at 0.5 mg/mL in hexane.

Cumulative **TGA Table** for **dodecathiol-coated AuNPs**

| Sample | Decomp. T1 (ºC) | Weight Loss 1 (%) | Decomp. T2 (ºC) | Weight Loss 2 (%) |
|---|---|---|---|---|
| 1 | 307.3 | 11.4 | - | - |
| 2 | 296.9 | 11.7 | - | - |
| 3 | 299.6 | 12.0 | - | - |
| Ave | 301 ± 5 | 11.7 ± 0.3 | - | - |

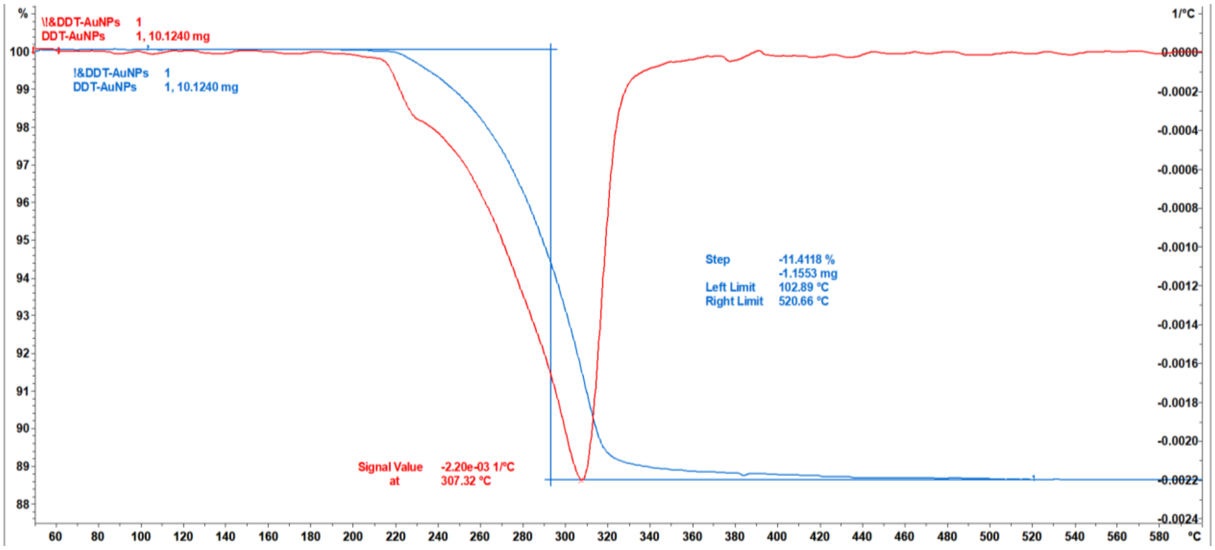


TGA of C12-S-AuNPs #1

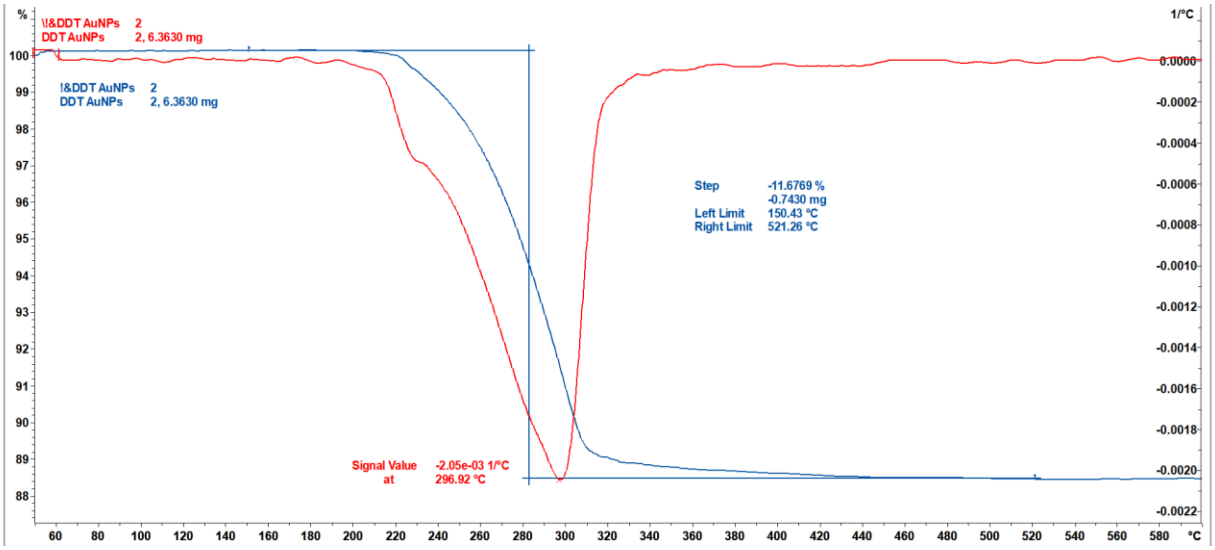


TGA of C12-S-AuNPs #2

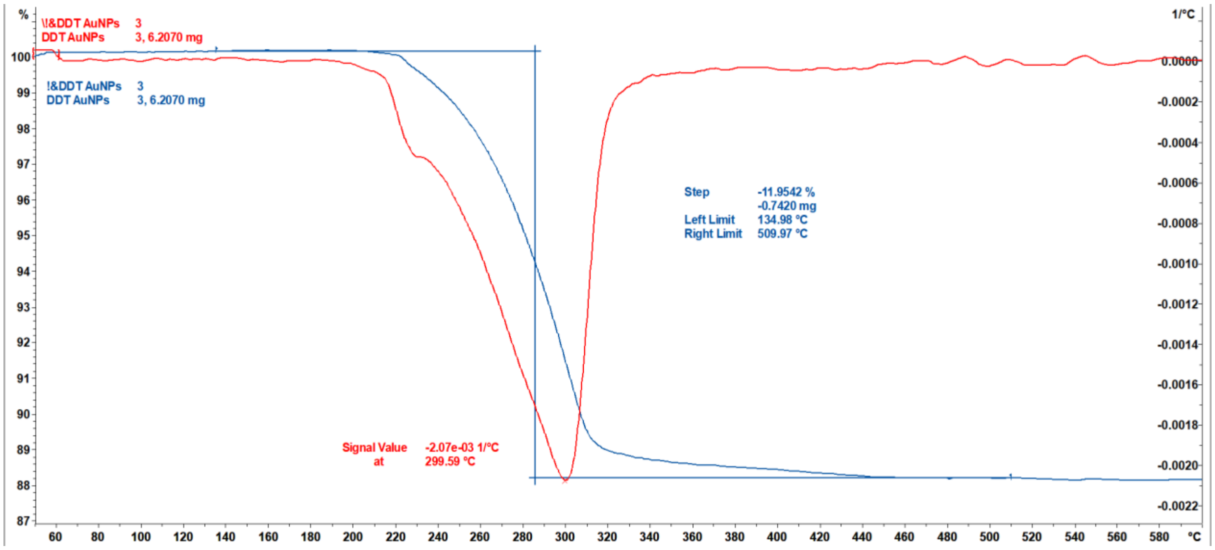


TGA of C12-S-AuNPs #3

**Figure S8**. TGA graphs of dodecathiol-coated AuNPs.

Cumulative **TGA Table** for **HOOC-PEG-SH**

| Sample | Decomp. T1 (ºC) | Weight Loss 1 (%) | Decomp. T2 (ºC) | Weight Loss 2 (%) |
|---|---|---|---|---|
| 1 | - | - | 416.1 | 96.8 |
| 2 | - | - | 415.8 | 96.6 |
| 3 | - | - | 415.2 | 96.4 |
| Ave | - | - | **416** ± 1 | **96.6** ± 0.2 |

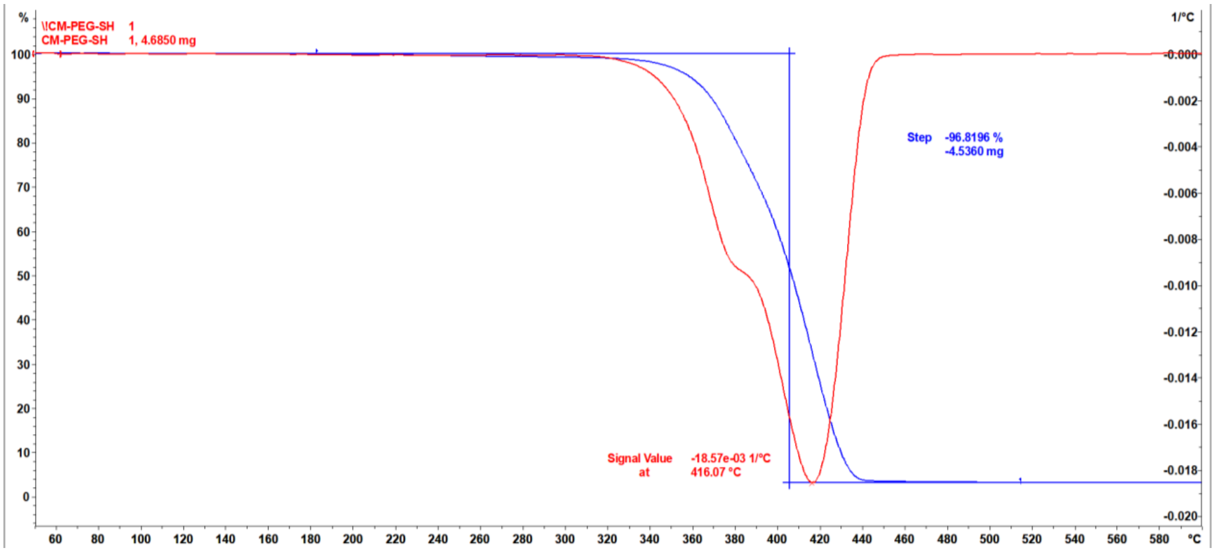


TGA of HOOC-PEG-SH #1

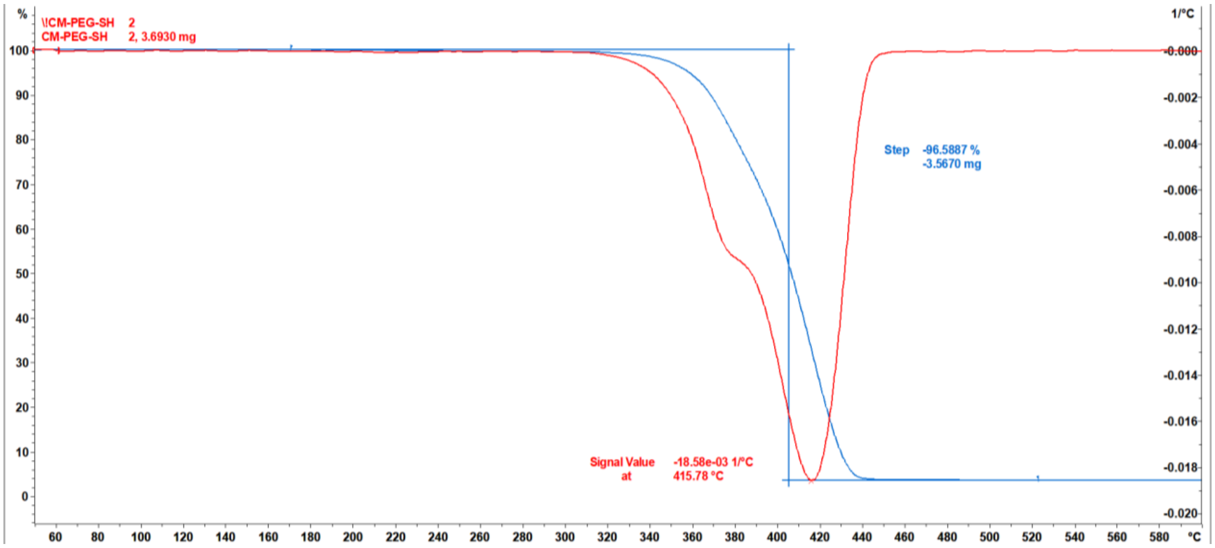


TGA of HOOC-PEG-SH #2

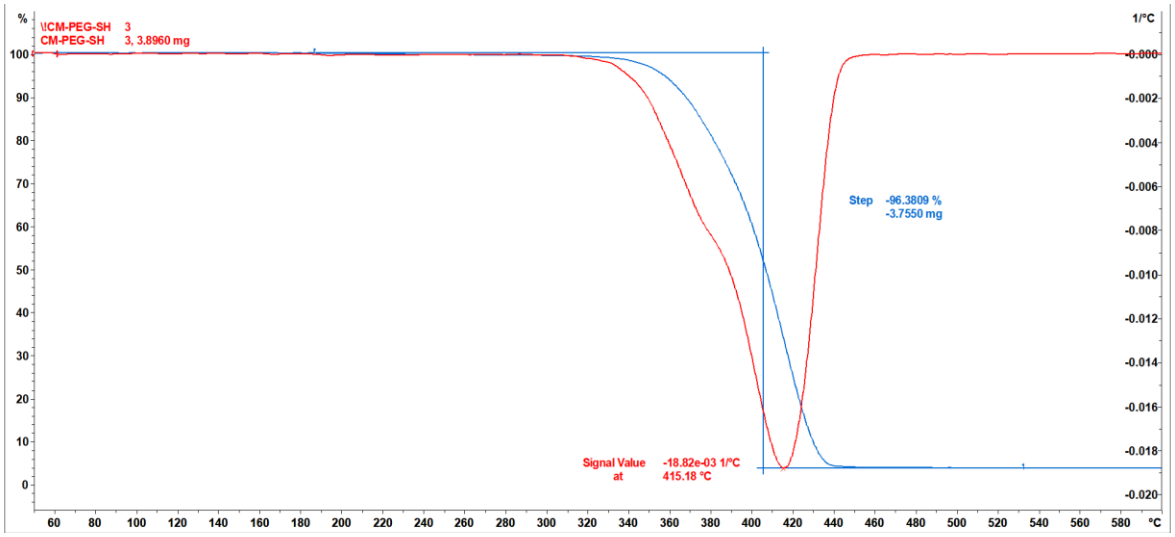


TGA of HOOC-PEG-SH #3

**Figure S9**. TGA graphs of **HOOC-PEG-SH**.

**TGA Table of *purified* HOOC-PEG-AuNPs**

| **Sample** | **Decomp. T1 (ºC)** | **Weight Loss 1 (%)** | **Decomp. T2 (ºC)** | **Weight Loss 2 (%)** |
|---|---|---|---|---|
| 1 | 277.9 | 5.9 | 404.5 | 33.6 |
| 2 | 282.4 | 6.2 | 400.6 | 35.1 |
| 3 | 276.1 | 5.8 | 398.8 | 34.8 |
| Ave | **279** ± 3 | **6.0** ± 0.2 | **401** ± 3 | **34.5** ± 0.8 |

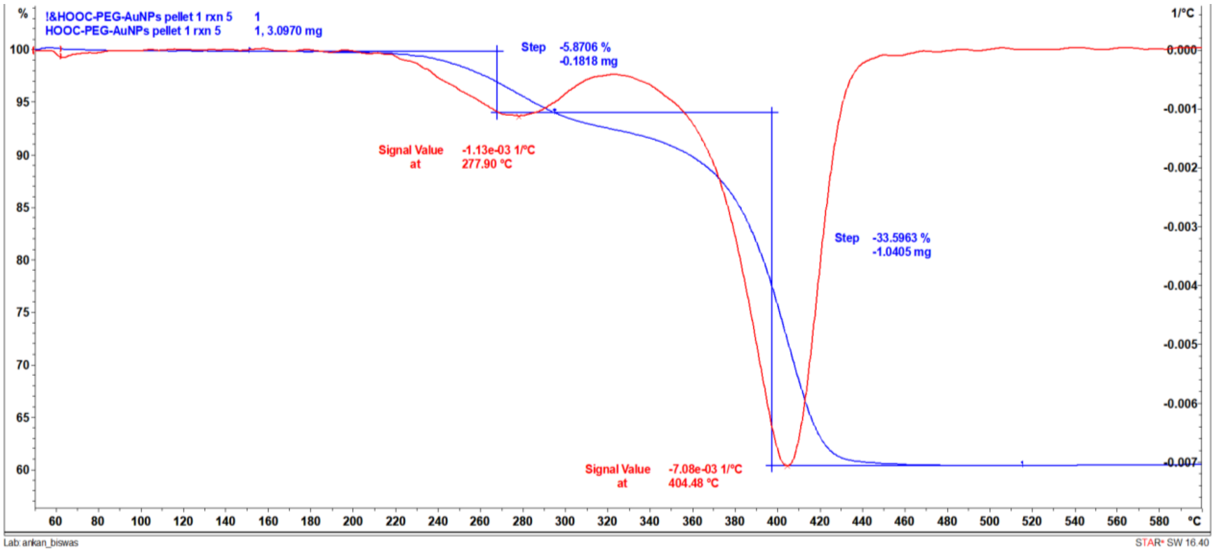


HOOC-PEG-AuNPs pellet #1

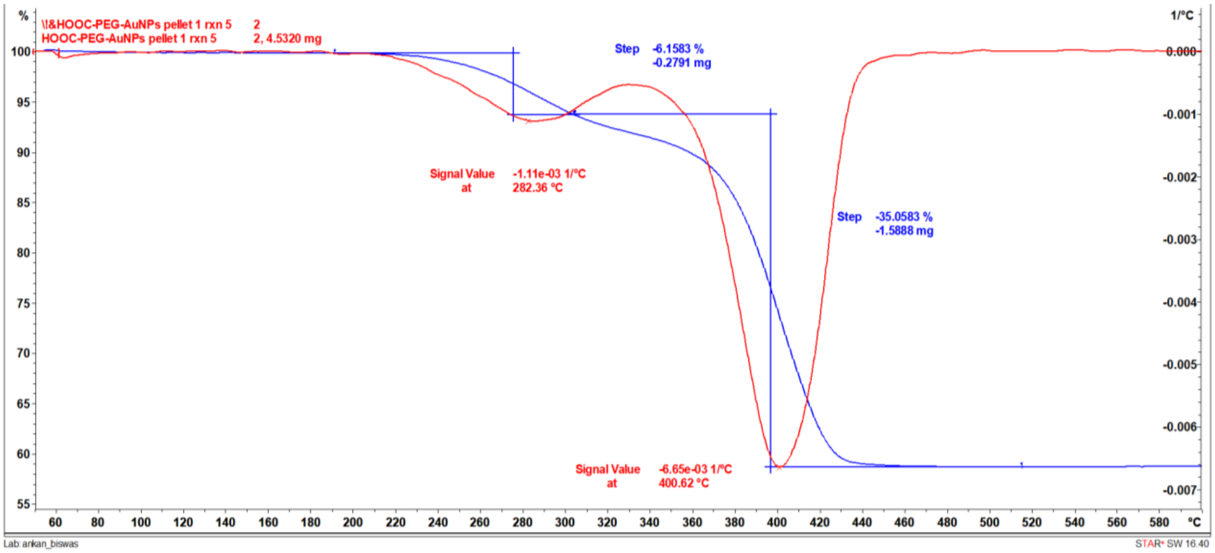


HOOC-PEG-AuNPs pellet #2

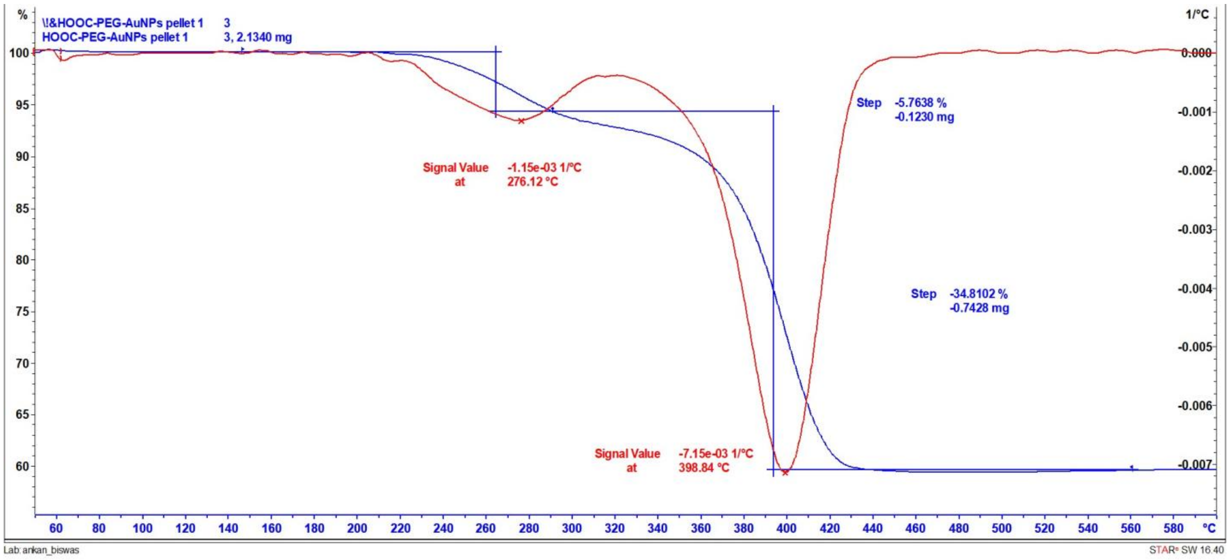


HOOC-PEG-AuNPs pellet #3

**Figure S10**. TGA graphs of **purified HOOC-PEG-AuNPs**.

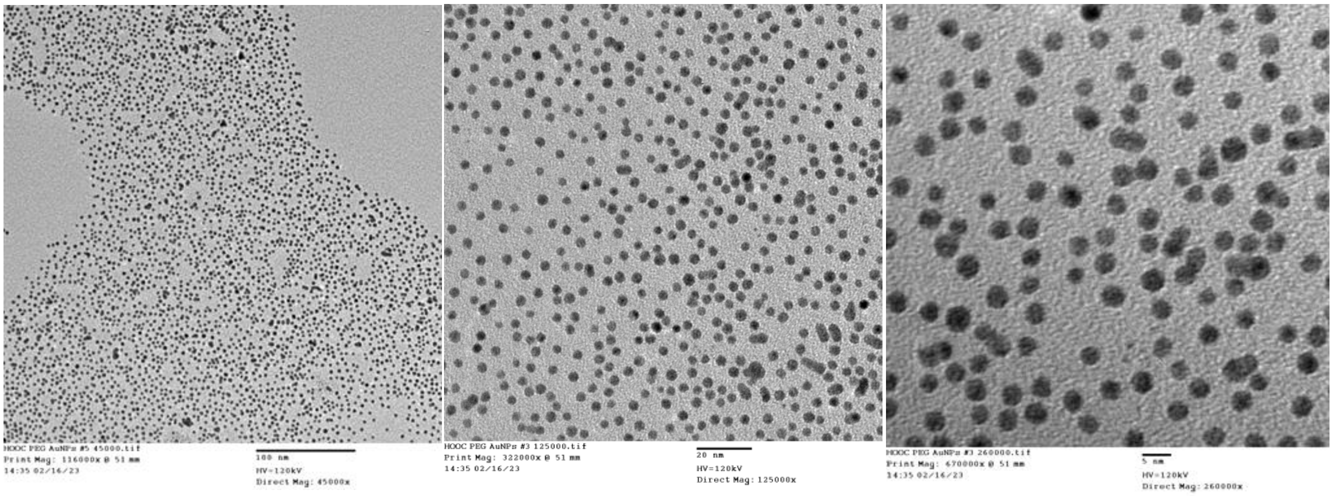


**Figure S11**. TEM images of **3.2** ± 0.4 **nm** HOOC-PEG-AuNPs at 0.5 mg/mL in water. The examined magnitudes (left to right) were 45000x, 125000x and 260000x.

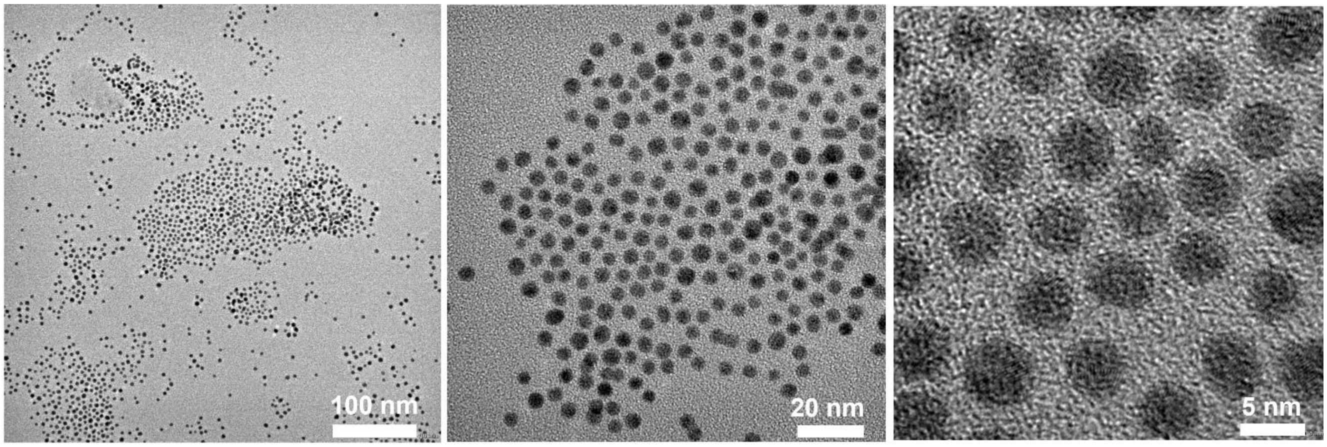


**Figure S12**. TEM images of **4.2** ± 0.6 **nm** *GAT1508*-PEG-AuNPs at 0.5 mg/mL in water at different magnifications (left to right).

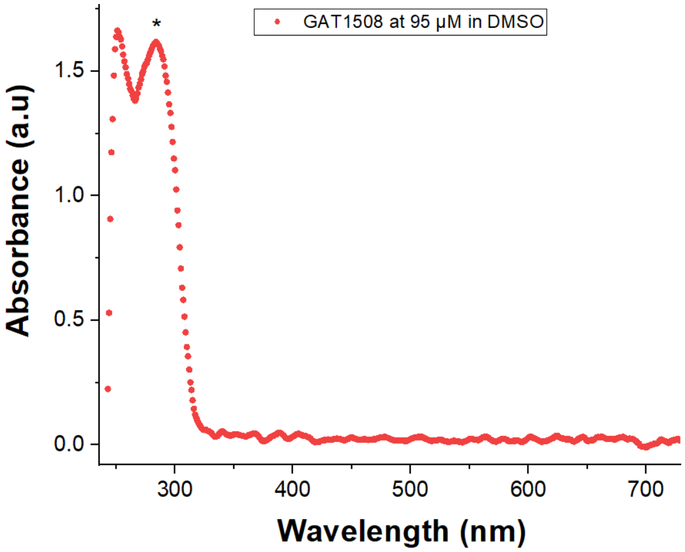


**Figure S13**. UV-Vis of *GAT1508* at 95 µM in DMSO.

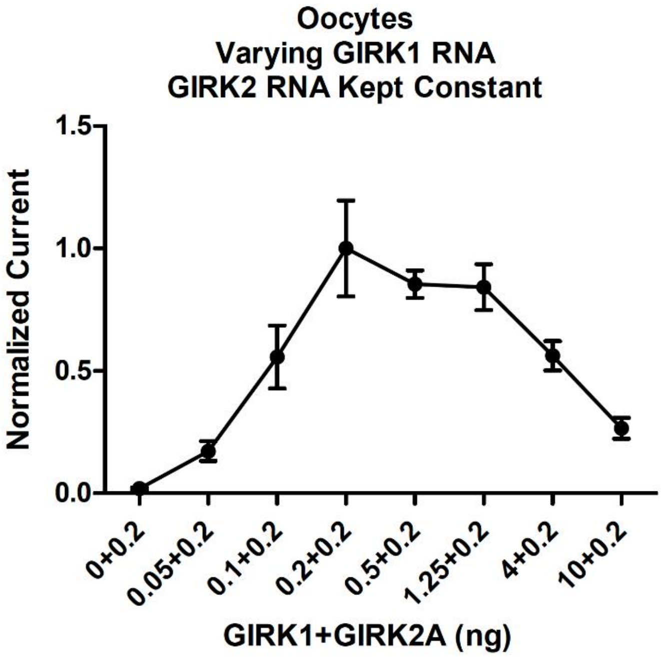


**Figure S14.** GIRK1/GIRK2 injected RNA ratios from 0.25:1 up to 50:1 can lead to the formation of transmembrane GIRK1/2 tetrameric units enabling ion channel-opening and flow of electric current.

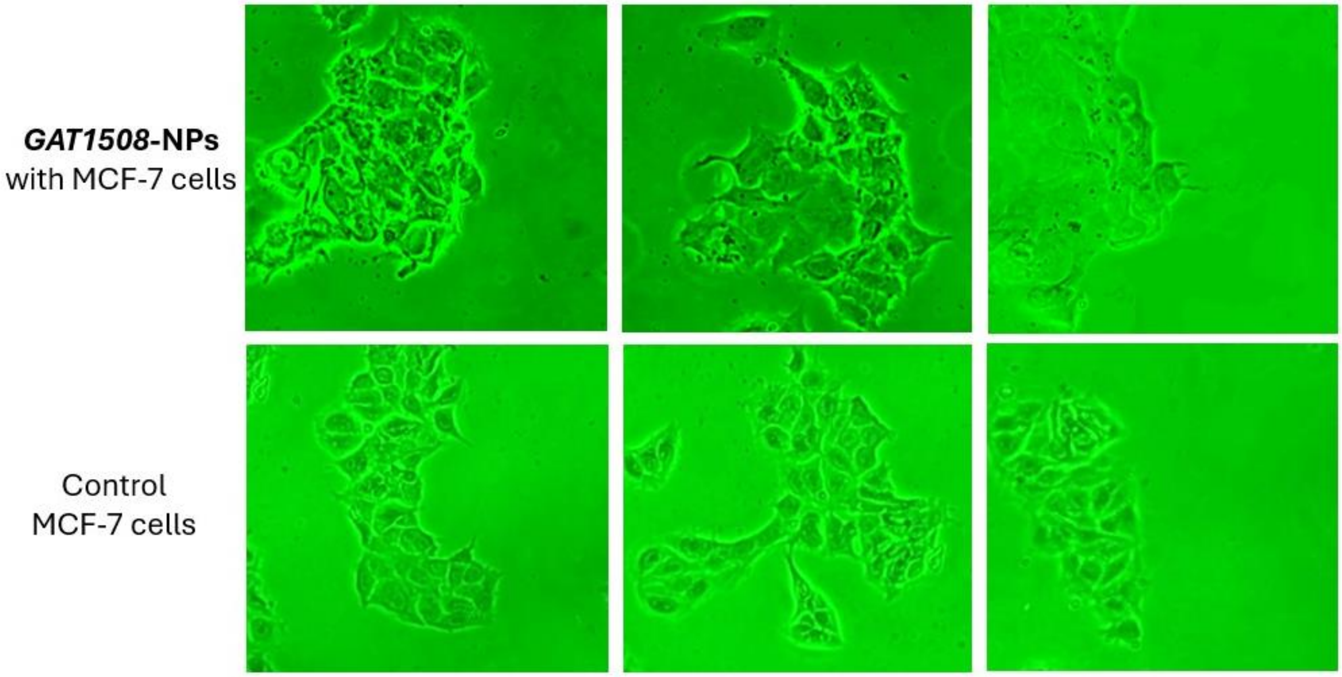


**Figure S15.** Optical microscope images of MCF-7 cells in the presence of GAT1508-coated NPs obtained at a different (basic) optical microscope with a cell phone camera.

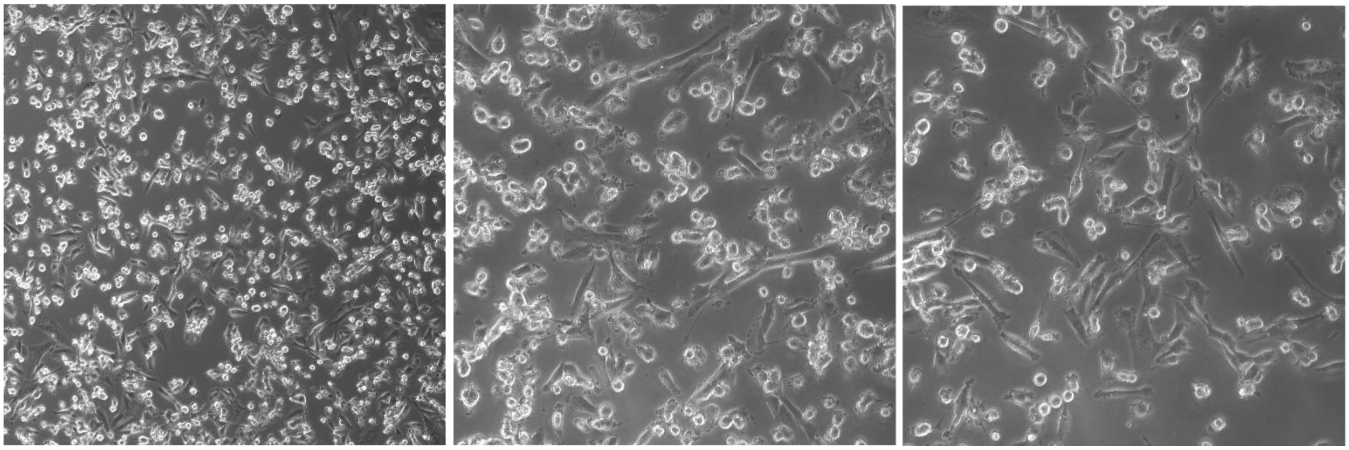

**Figure S16.** Optical microscope images of triple (-) MDA-MB-231 cells that do not express GIRK1/2 in after incubation with *GAT1508*-coated NPs showing absence of interaction.

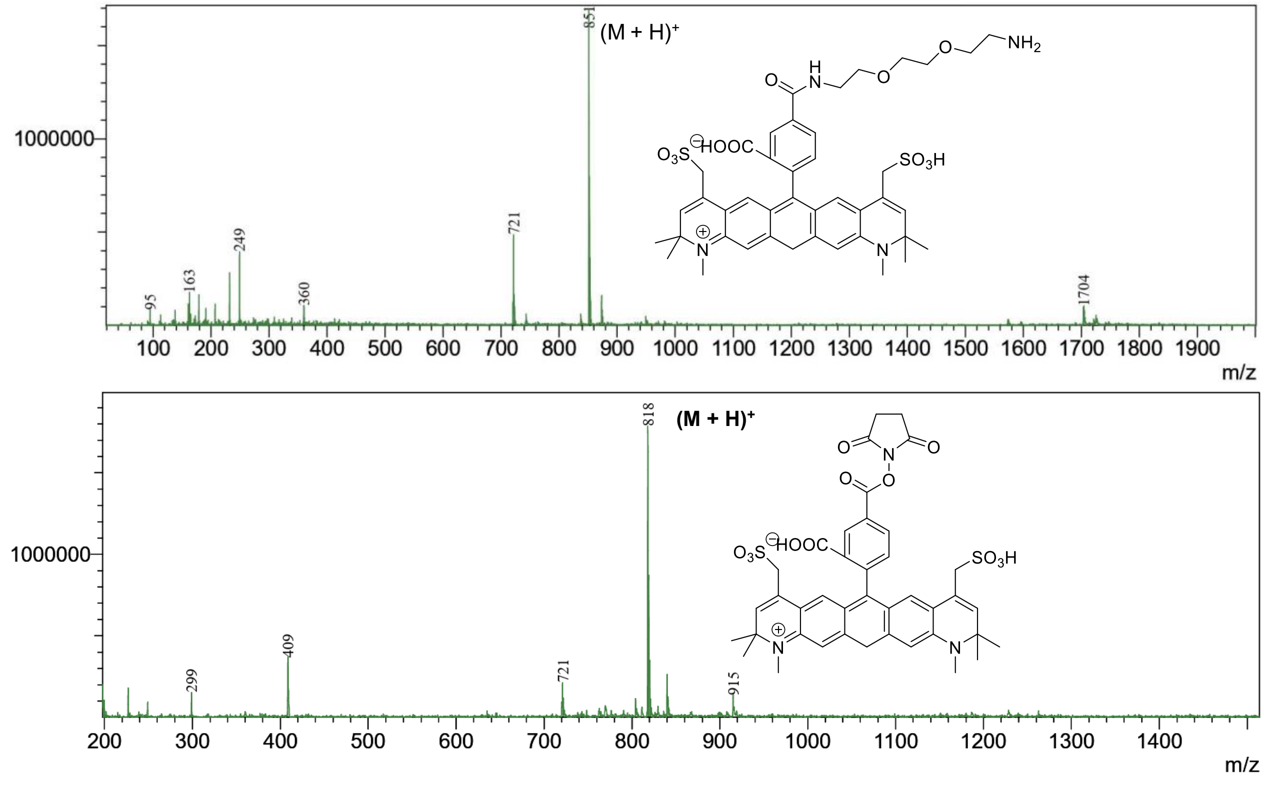


**Figure S17.** LC-MS spectra of (a) Alexa Fluor 570-$EG_2$-$NH_2$ (MW = 851 g/mol) and (b) starting material Alexa Fluor 594 NHS ester (MW = 818 g/mol).
